# In-Depth Characterization of Apoptosis N-terminome Reveals a Link Between Caspase-3 Cleavage and Post-Translational N-terminal Acetylation

**DOI:** 10.1101/2022.09.19.508487

**Authors:** Rawad Hanna, Andrey Rozenberg, Layla Saied, Daniel Ben-Yosef, Tali Lavy, Oded Kleifeld

## Abstract

The N-termini of proteins contain information about their biochemical properties and functions. These N-termini can be processed by proteases and can undergo other co- or post-translational modifications. We have developed LATE (LysN Amino Terminal Enrichment), a method that uses selective chemical derivatization of α-amines to isolate the N-terminal peptides, in order to improve N-terminome identification in conjunction with other enrichment strategies. We applied LATE alongside another N-terminomic method to study caspase-3 mediated proteolysis both *in vitro* and during apoptosis in cells. This has enabled us to identify many unreported caspase-3 cleavages, some of which cannot be identified by other methods. Moreover, we have found direct evidence that neo-N-termini generated by caspase-3 cleavage can be further modified by Nt-acetylation. Some of these neo-Nt-acetylation events occur in the early phase of the apoptotic process and may have a role in translation inhibition. This has provided a comprehensive overview of the caspase-3 degradome and has uncovered previously unrecognized crosstalk between post-translational Nt-acetylation and caspase proteolytic pathways.

## Introduction

Proteases used to be described as degradative enzymes that disassemble unwanted or damaged proteins to generate amino acids for the cell (Lopez-Otin & Overall, 2002; López-Otín & Bond, 2008). Many proteases, however, are very precise and specific enzymes, and their activity produces new polypeptides for different purposes (Lopez-Otin & Overall, 2002; Schlage *et al*, 2017). Such specific proteolytic cleavages are irreversible post-translational modifications. They are crucial for cellular processes and therefore tightly regulated (Marino *et al*, 2015; Sukharev *et al*, 1997; Tafani *et al*, 2002; Zeeuwen, 2004). Thus, characterization of protease activity and substrate specificity are essential for completing the functional annotation of any proteome.

Mass-spectrometry (MS) based proteomics is one of the most commonly used methods for system-wide protein studies. Despite this, there are plentiful challenges to overcome when MS is used to identify and quantify proteolytic substrates (Rogers & Overall, 2013). These challenges predominantly stem from the low abundance of proteolytic fragments, which can easily be masked by other peptides. Moreover, background *in vivo* proteolysis events and the transient state of protein synthesis and degradation also contribute to the system complexity when studying a specific protease activity or treatment-directed proteolysis. Over the past decades, various approaches have been developed for reliable detection of proteolytic fragments by separating them from the background proteome (Luo *et al*, 2019). N-terminomics methods are used to isolate and characterize the N-terminal fragment of every protein (N-terminome), usually by utilizing a series of chemical reactions to manipulate the desired subgroup. Negative selection methods such as combined fractional diagonal chromatography (COFRADIC) (Gevaert *et al*, 2002), terminal amine isotopic labeling of substrates (N-TAILS) (Kleifeld *et al*, 2010), and hydrophobic tagging-assisted N-termini enrichment (HYTANE) (Chen *et al*, 2016) are based on the depletion of internal peptides to enhance the identification of proteolytic products N-terminus (neo N-termini) and original N-termini, including naturally modified ones. Positive selection techniques are based on the enrichment of protein fragments containing unblocked (or free) N-termini after tagging them with biotin (Mahrus *et al*, 2008), and therefore cannot be used to study the majority of eukaryotic N-terminal peptides that are naturally blocked (Griswold *et al*, 2019; Mahrus *et al*, 2008).

Each method has advantages and limitations weighted by its reliability, sensitivity, availability, cost, and time consumption. For example, COFRADIC can be applied to study the N-terminome with low sequence biases using commercially available reagents, but it requires a long analysis time and therefore extensive instrumental usage which can make it difficult to analyze many samples at once (Gevaert *et al*, 2002; Luo *et al*, 2019). N-TAILS, on the other hand, is simpler and more straightforward but requires the use of a specific polymer (Kleifeld *et al*, 2010, 2011). A recently developed method based on hydrophobic tagging akin to HYTANE, termed high-efficiency undecanal-based N-termini enrichment (HUNTER) (Weng *et al*, 2019) utilizes cheaper reagents and has shown to be useful for low protein amount samples. Most of the negative selection N-terminomics approaches require primary amine labeling and thus involve blocking of lysine ε-amines. As a result, these methods cannot use proteases with lysine cleavage specificity, which significantly restricts their application in annotation of the N-terminome.

The N-terminus of the nascent polypeptide chain as translated from its open reading frame (ORF) initiation site is the first to undergo modifications. Such modifications include N-terminal methionine excision and acetylation which usually take place co-translationally when the protein is still short and bound to the ribosome (Giglione *et al*, 2015). For the majority of proteins, the N-terminal methionine is removed by methionine aminopeptidases (Jonckheere *et al*, 2018). Nt-acetylation is catalyzed by N-terminal acetyltransferases which utilize acetyl-CoA as the donor for the transfer of the activated acetyl moiety to the N-terminus of the protein (Ree *et al*, 2018). It is estimated that ∼80% of human proteins are modified by N-terminal acetylation (Nt-acetylation). By altering the N-terminal charge and hydrophobicity of a protein, Nt-acetylation can affect a range of its properties including stability, folding, protein-protein interactions, and subcellular targeting (Ree *et al*, 2018).

To improve the coverage of the N-terminome, we developed LysN Amino Terminal Enrichment (LATE), an N-terminal enrichment strategy based on LysN digestion and specific N-terminal α-amine isotopic labeling (Figure 1B). We employed LATE in parallel to the HYTANE N-terminomics method to study the central cellular apoptosis protease, caspase-3. By applying this approach to a set of cell-based apoptosis experiments and controlled *in vitro* studies, we identified many reported as well as novel caspase-3 cleavages. With this comprehensive mapping of N-terminal peptides, we demonstrate how Nt-acetylation affects the ORF N-terminal susceptibility to cleavage by caspase-3 and find that caspase-3 cleavage can generate new sites for post-translational N-terminal acetylation. We also show that these Nt-acetylation (neo-Nt-acetylation) events occur in the early stages of apoptosis. This is the first demonstration of a link between post-translational neo-Nt-acetylation and caspase proteolytic pathways.

**Figure 1.**
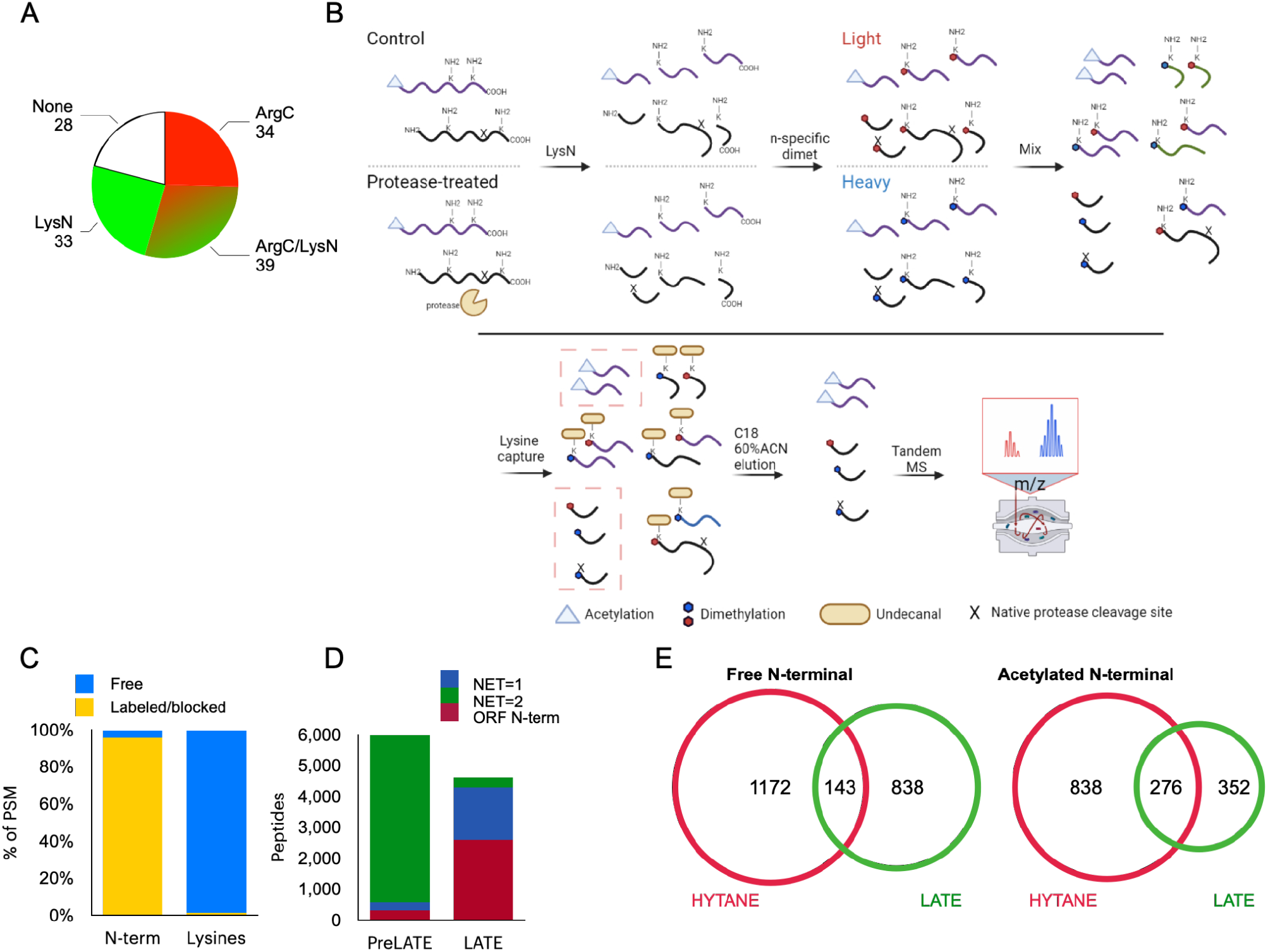
LATE workflow for improved N-terminome coverage. **A**. *In silico* analysis of the “Identifiable” and “Not identifiable” putative caspase-3 DEVD↓X in the human proteome following digestion by trypsin with ArgC-like specificity (red) and LysN (green) **B**. LysN Amino Terminal Enrichment (LATE) workflow for the comparison of two conditions. Extracted proteins of each sample are digested with LysN, and the resulting peptides undergo N-terminal-specific isotopic dimethylation. The samples are then mixed and subject to hydrophobic tagging of lysines which allows for their removal and for the enrichment of the N-terminal peptides **C**. LATE N-terminal dimethylation labeling specificity following LysN digestion. Over 95% of peptides’ N-terminal amines were labeled by dimethylation or naturally blocked by acetylation while less than 5% of the lysine side-chain amines were labeled **D**. Peptide-spectrum match (PSM) number of LysN internal peptides (green) versus the PSM number of ORF (burgundy) and neo N-terminal peptides (blue) before (PreLATE) and after LATE **E**. Comparison of Human proteins with free (left) and acetylated (right) N-terminal peptides identified by HYTANE (red) and LATE (green).

## Materials and Methods

### Chemicals and reagents

All chemicals and reagents were purchased from Sigma-Aldrich unless specified otherwise. Undecanal was purchased from TCI, Acetonitrile was purchased from JT Baker, formic acid was purchased from Biolabs ltd, Guanidium-HCl and HEPES were purchased from the Spectrum, and ethanol was purchased from LiChorsolv. HPLC-grade water was purchased from Thermo. All solvents used are MS-grade.

### Plasmids

The construct for *E*.*coli* expression of WT caspase-3 (pET23b-Casp3-His) was obtained from Addgene (plasmid 11821). The construct for mammalian expression of caspase-3/GFP was a gift from Dr. Yaron Fuchs Lab (Technion).

### *E. coli* Proteome Preparation

*E. coli* DH5α cells were grown in LB until an optical density of OD_600nm_=0.5. Cells were harvested by centrifugation at 6000 g for 10 min at 4°C and washed twice with 50 mM PBS pH 7.4. The cells pellet was resuspended in 50 mM HEPES, 8M Gu-HCl pH 7.5, and lysed by heating to 95°C for 10 min. The lysate was clarified by centrifugation at 15000 g for 10 min at 4°C, and the supernatant was collected.

### Caspase-3 expression and purification

The expression construct of pET23b-Casp3-His6 was extracted from DH5α bacterial culture purchased from Addgene (Plasmid #11821). BL21-CodonPlus (DE3)-RIPL competent cells were transformed with the expression construct and plated on LB-agar supplemented with ampicillin (100 µg/ml) and chloramphenicol (34 µg/ml). A single colony of the transformants was inoculated into a 5 ml LB supplemented with the above antibiotics and cultured at 37°C and 200 rpm shaking, overnight. On the next morning, the overnight culture was inoculated into 50 ml LB supplemented with the above antibiotics and grown for 3 hours at 37°C and 200 rpm shaking. The culture was transferred to 1L of LB in a baffled flask and cultured. At OD_600nm_=0.6, the temperature was lowered to 17°C and expression was induced by the addition of IPTG (1 mM). Expression was terminated after 16 hours by harvesting the cells (centrifugation for 20 min at 4000g at 4°). The harvested cells were resuspended in 50 mL ice-cold buffer A (200 mM NaCl, 20 mM Tris-HCl, pH 7.5) and stored at -80°C. The cells were lysed by three freeze-thaw cycles (liquid nitrogen/RT) followed by disruption of the cells using a cell disrupter (OS Model, Pressure Biosciences). The lysate was clarified by centrifugation (20,000 g, 4°C, 20 minutes) and the clear lysate was purified on HisTrap HP 1 ml affinity column (Cytiva) by FPLC system (NGC™, BioRad). The protein was eluted from the column using a linear gradient (Elution buffer: 0-1M imidazole, 200 mM NaCl, 20 mM TRIS-HCl pH 7.5). The fractions were analyzed by SDS-PAGE. Based on SDS-PAGE, fractions containing caspase-3 were pooled together and dialyzed against 20 mM HEPES 7.5, and 80 mM NaCl. Aliquots of the protein were stored at -80°C.

### Cell culture

Human colorectal carcinoma cells (HCT116) were grown in RPMI supplemented with 10% fetal calf serum (FCS), 2mM L-Glutamine, and 1% Pen-Strep solution. Immortalized aneuploid human keratinocyte cells (HaCaT) were grown in DMEM supplemented with 10% FCS, 2mM L-glutamine, and 1% Pen-Strep solution, in 10 cm plates. When the cells reached 80% confluence, the medium was discarded, and the cells were washed three times on the plate with 10 ml PBS before detachment or lysis.

For cell-based caspase-3 experiments, HCT116 were grown in a 10 cm plate until they reached 40% confluence, then the medium was changed to starvation medium (RPMI 2mM L-glutamine, 1% Pen-Strep solution without serum) for 2 hours. The transfection mix was prepared by mixing 15 µl Polyjet (SignaGen) with 6 µg of caspase-3/GFP bicistronic vector each prepared in 250 µl starvation media, the mix was left for 10 min at room temperature before adding it to the starved cells. Eight hours following transfection the media was changed back to RPMI supplemented with 10% FCS, 2 mM L-glutamine, 1% Pen-Strep solution, and GFP fluorescence was observed to estimate the efficiency. Next, the cells were split and grown to 80% confluence before without sorting.

For time-course studies HCT116 cells were grown in a 6-well plate until they reached 60% confluence, then cells were washed twice before it was changed to fresh RPMI media (as above) with 150µM ABT-199 or DMSO (control). Cells were collected from the ABT-treated and control cells after 0.5, 1 and 2 hours after ABT-199 (or DMSO) addition.

### Cell viability assay

Ten thousand HCT-116 transfected with caspase-3/GFP bicistronic vector or non-transfected cells were seeded in 96-well plates and grown for 24 hours in RPMI media (as described above). Then the cells were treated (in triplicates) with different concentrations of ABT-199 (0, 10, 50, 100, 150 µM). At each of the specified time points cell viability was measured using XTT-based Cell Proliferation Kit (Sartorius), according to the manufacturer’s protocol.

### Cell lysis and protein extraction

For extraction under denaturative conditions, 1ml of 95°C pre-heated 50mM HEPES, 8M Gu-HCl pH 8.0 was added to each plate, and cells were scraped down and incubated 5 min at 95°C. Next, the cell lysates were sonicated using UP200St with VialTweeter (Hielscher, Germany) at max amplitude, 70% cycle time for 5 minutes. The sonicated lysates were centrifuged at 15000g, 10 min, 4°C, and the cleared supernatants were collected. For extraction under native conditions, cells were detached following 5 minutes of treatment with trypsin-EDTA and collected by centrifugation (5000g, 5min, 4°C). Cell pellets were washed three times with PBS, and resuspended in 0.5 mL of 150mM NaCl, 20mM HEPES pH 7.5, and lysed by 3 freeze-thaw cycles of 5 minutes in liquid nitrogen followed by 20 minutes on ice. The lysate was clarified by centrifugation at 15000g for 10 min at 4°C. Protein concentration was determined by BCA assay (Smith *et al*, 1985).

### *In vitro* caspase experiments

Clear lysate extracted under native conditions was split into two aliquots. One was left untreated (control) while the other was mixed with recombinant human caspase-3 in a 1:20 w/w ratio. Both samples were kept at 37°C with gentle mixing. After 1, 6, and 18 hours, a fraction of 200 ug protein was taken from the control and caspase-3-treated samples into a clean tube containing an equal volume of 20mM HEPES,8M Gu-HCl pH 7.5 and incubated at 95°C for 5 minutes. For checking *in vitro* Nt-acetylation, the clear lysate was split into two aliquots. One was dialysed by 3 cycles of concentration and 10X dilution at 4°C in ice-cold 20mM HEPES pH7.8, 1mM DTT,1mM EDTA, 100mM NaCl using 3k cut-off Amicon® Ultra 2 mL Centrifugal Filters (Merck). Equal amounts of protein from both dialyzed and undialyzed samples were treated with caspase-3 as described above. Aliquots were taken from each sample after 0.5, 6 and 18 hours. Nt-acetylation samples were subject only to HYTANE.

### Cell-based caspase experiments

HCT-116 cells transfected caspase-3/GFP vector were detached with trypsin-EDTA, quenched by an equal volume of media, and centrifuged at 1500g for 5min. Then the cells were resuspended in 1ml RPMI supplemented with 10% FCS, 2mM L-Glutamine, 1% Pen-Strep solution, and transferred to a 5mL polystyrene round-bottom tube with a cell-strainer cap (Falcon). Unstained negative control cells were also prepared in the same way for population determination during sorting. Sorting was done using BD FACS-Aria™ (BD Bioscience) against GFP fluorescence. Between 1.5 and 2 million cells of GFP-positive (high caspase-3) and negative populations were collected per biological repeat. After sorting, samples were centrifuged at 1500g for 5min, re-suspended in 4 ml rich media (RPMI, 20% FCS, 1% Pen-Strep, 2mM L-Glutamine), and seeded in a 6 cm plate. Cells were grown for another day while the media refreshed after 12 hours. Lastly, samples were treated with BCL-2 inhibitor ABT-199 at a final concentration of 150µM to induce caspase-3 activation for 2 hours, prior to harvesting the cells.

### LysN Amino Terminal Enrichment (LATE)

One hundred ug protein was re-suspended in 50mM HEPES, 4M Gu-HCl pH 7.5and reduced with 5mM DTT at 65°C for 30 minutes followed by alkylation with 12.5mM chloroacetamide (CAA) for additional 20 minutes at room temperature in the dark. CAA leftovers were quenched with another dose of DTT, and the sample was diluted 1 to 4 with 20mM HEPES pH 8 and the pH was further adjusted to 8.0 if needed using 1N NaOH. Next, LysN (Promega) was added at 1:100 w/w and the samples were digested for 12h at 37°C. Digestion was terminated by adding a 6% final volume of glacial acetic acid. If required additional acetic acid was added until the pH reached the range of 2.6-2.8. N-terminal alpha amines specific labeling was done by adding 40mM (final) of C^13^D_2_ (heavy) or C^12^H_2_ (light) formaldehyde and 20mM (final) of sodium cyanoborohydride at 37°C for 10min. Heavy and light labeled samples were quenched by addition of glycine (final concentration 100mM) dissolved in 2% acetic acid and incubation at room temperature for 10min. Next heavy- and light-labeled samples were mixed and desalted using OASIS-HLB 100mg cartridge (Waters). Peptides were eluted with 60% ACN, and 0.1% FA and dried to completeness using CentriVap Benchtop Vacuum Concentrator (Labconco). Next, samples were resuspended in 100µl of 100mM HEPES pH 7.0 with undecanal at 50:1 unedecanal:peptide w/w ratio. Undecenal was dissolved in ethanol to a final concentration of 20mg/ml. The undecanal labeling was carried out at 50°C for 2 hours.

Undecanal-labeled samples were centrifuged at 20000g for 5 minutes at RT and the supernatants were transferred to a new tube and dried to completeness using CentriVap Benchtop Vacuum Concentrator (Labconco, USA). Dried samples were dissolved in 2% ACN, and 0.1% FA and desalted with OASIS-HLB 100mg cartridge (Waters). Undecanal leftovers and undecanal-modified peptides should remain bound to the column after elution with 60% ACN. Samples were dried and resuspended in 2% ACN, and 0.1% FA prior to LC-MS analysis.

### Hydrophobic tagging-assisted N-termini enrichment (HYTANE)

One hundred ug protein of each of the compared samples were resuspended in 100mM HEPES, 4M Gu-HCl pH 7.5, and reduced with 5mM DTT at 65°C for 30 minutes followed by alkylation using 12.5mM CAA for additional 20 minutes at room temperature in the dark. CAA leftovers were quenched with another dose of DTT. Each sample was labeled by adding 40mM (final) of either C^13^D_2_ (heavy) or C^12^H_2_ (light) formaldehyde and 20mM (final) of sodium cyanoborohydride and incubating at 37°C overnight followed by quenching with 100mM glycine (final) for 1 hour at 37°C. Next, heavy and light-labeled samples were mixed and diluted 1 to 4 using 20mM HEPES pH 8.0. The mixed samples were digested with sequencing grade trypsin (Promega) in a 1:100 w/w ratio, at 37°C overnight. The tryptic digestion was quenched by the addition of FA to a final concentration of 1%. The combined sample was desalted using OASIS-HLP and dried completely in a speedvac. Undecanal-based enrichment was done as described for LATE.

### LC-MS

Desalted samples were subjected to LC-MS analysis using a Q Exactive Orbitrap Plus mass spectrometer Q Exactive Orbitrap HF coupled to Easy-nanoLC 1000 capillary HPLC. The peptides were resolved by reversed-phase using a homemade 30 cm long 75µm-diameter capillary, packed with 3.5-µm silica using ReproSil-Pur C18-AQ resin (Dr. Maisch GmbH). Peptides were eluted using a linear 120-minute gradient of 5–28% acetonitrile with 0.1% formic acid, followed by a 15-minute gradient of 28–95% acetonitrile with 0.1% formic acid, and a 15-minute wash of 95% acetonitrile with 0.1% formic acid (at flow rates of 0.15 μl/min). MS was performed in positive mode using an m/z range of 300–1800 in a data-dependent acquisition mode, a resolution of 70,000 for MS1, and a resolution of 17,500 for MS2; repetitively full MS scans were followed by high energy collisional dissociation (HCD) of the 10 or 20 most dominant ions selected from the first MS scan for Q-Exactive Plus and Q-Exactive HF respectively.

### Data analysis

Data-dependent acquisition experiments were analyzed using Trans Proteomic Pipeline (version 6.1 or 5.2) (Deutsch *et al*, 2010). Searches were done using COMET (version 2020_01 rev 1 or 2018_01 rev 1) (Eng *et al*, 2013). Searches for acetylated, dimethylated, and methylated N-termini were done separately using the search parameters listed in Table S11. Comet results were further validated by PeptideProphet (Keller *et al*, 2002) at a false discovery rate of less than 1%. XPRESS was used for relative peptide quantification (Han *et al*, 2001). Mass tolerance for quantification was 20 ppm with The minimum number of chromatogram points needed for quantitation was set to 3, and the number of isotopic peaks to the sum was set to 0. Searches were conducted against *Homo sapiens* proteome containing 73248 sequences (ID:9606, downloaded from Uniprot in June 2019) and supplemented with cRAP sequences downloaded from ftp://ftp.thegpm.org/fasta/cRAP. Bioinformatic analysis of the N-terminome identifications was carried out using in-house python scripts (available for download from the github repository https://github.com/OKLAB2016/peptide-matcher and https://github.com/OKLAB2016/parse-and-unite) and based on Uniprot annotations (The UniProt Consortium, 2021). These scripts were used to combine peptide identification into unified list, aggregate peptide ratios based on the sum of peak areas of different the PSM of the same peptide, map peptide location with the protein sequence and provide secondary structure and relative solvent accessibility for the peptides and its flanking amino acids. Differential amino acid usage logos were created with dagLogo v. 1.32.0 (Ou *et al*, 2020). Logos were plotted as the percent difference of the amino acid frequency at each site against the background of the Swiss-Port human proteome. Boxplots were created using BoxPlotR (Spitzer *et al*, 2014). GO term enrichment analysis was performed with topGO v. 2.46.0 (Alexa & Rahnenfuhrer, 2022) and the results were visualized by plotting terms significantly enriched with respect to the whole human proteome (FDR≤0.01) in semantic coordinates obtained with rrvgo v. 1.6.0 (Sayols, 2022).

### Experimental Design and Statistical Rationale

Cell-based proteomic analyses were done using triplicate cell cultures (biological replicates). Biological replicates were individual cultures grown separately and prepared separately. Once these cultures reached appropriate confluence levels, they were treated as described. *In vitro* proteomics analyses were done using a protein extract from one cell culture (one biological repeat). This extract was aliquoted, and each aliquot was subject to different treatments such as +/-dialysis and/or +/-incubation with caspase-3. These different treatments were sampled at different time points. The quantitative proteomics analysis was based on isotopic labeling. The abundance ratios of each sample (MS run) were normalized against the median ratio and then quantitative data from identical peptide samples were aggregated together by summation. Significant ratio changes were defined based on a fixed threshold as described before (Kleifeld *et al*, 2010). All data will be publicly available and respective repositories.

## Results

Most proteomics studies aimed at studying proteolytic processing and modification of protein N-termini rely on negative selection methods. Many of these N-terminomics methods are based on blocking/labeling of lysine residues which prevents the use of proteases with lysine cleavage specificity, such that even though they utilize trypsin digestion, the resulting peptides are only those derived from cleavage at arginine residues (thus resembling ArgC digestion). This restriction limits the identification potential considerably; for example, identification of ORF N-termini is possible for only less than 50% of the human proteome (Figure S1). While repeating the N-terminomics experiments with proteases other than trypsin can improve the coverage, there are still several thousand protein ORF N-termini that can only be identified by a lysine-specific protease (Figure S1). A major use of N-terminomics methods is for the identification of proteolytic cleavage sites. For example, in the human proteome, there are 134 putative cleavages of caspase-3 the apoptosis executioner protease whose canonical cleavage motif is DEVD↓X (Walsh *et al*, 2008). Of these, only 73 (54%) can be identified by N-terminomics methods that are based only on cleavages at arginine residues. The number of identified caspase-3 cleavage sites can be considerably expanded to 106 (∼80%) by also using peptides generated from cleavage at lysine residues following LysN digestion (Figure 1A). Similar expansion in the number of identifications was obtained when a less restrictive cleavage motif was used. Aside from the sequence coverage, it was also shown in previous studies that peptides with a basic residue closer to the N-termini or the C-termini have different chromatographic characteristics, which contributes to the peptide diversity between LysN and trypsin (ArgC-like) peptides (Tsiatsiani *et al*, 2016). Based on these considerations, we developed LATE, a new method utilizing LysN digestion followed by specific N-terminal α-amines blocking while minimizing lysine modification, then exploiting the free lysine ε-amine to deplete internal peptides (Figure 1B). After protein extraction and digestion, α-amines are labeled using reductive dimethylation in acidic buffer (Qin *et al*, 2012) and the compared samples are mixed. Internal peptides containing lysine residues are depleted by adding undecanal that reacts with lysine ε-amine and aid to separate them on a C18 column based on the added hydrophobicity (Chen *et al*, 2016; Weng *et al*, 2019).

A key element in the efficiency of LATE is the specific dimethylation of peptides’ N-termini (α-amines) without notable labeling of lysine residues (ε-amines). The conditions for this step were optimized following recent reports (Qin *et al*, 2012; Koehler *et al*, 2013). LATE labeling efficiency was tested on the pre-enriched sample where we observed labeling of free N-terminal α-amines with >95% efficiency, while only <5% of lysine side chain amines were dimethylated (Figure 1C and Table S1). Internal peptides that have Number of Enzymatic Termini (NET) = 2, those being full LysN peptides, dominated the pre-enrichment sample with more than 80% (Figure 1D) accompanied by a low percentage of ORF N-terminal (i.e. that begin at positions 1 or 2 of the predicted ORF) and neo-N-terminal peptides. Around 10-fold enrichment of ORF N-terminal and neo-N-terminal was achieved after applying LATE, with the incidence of internal peptides decreasing to less than 10%. The same enrichment trend was observed in our HYTANE sample (Figure S2), similarly to the previous reports (Chen *et al*, 2016; Weng *et al*, 2019). As predicted, application of LATE allowed identification of a significant number of dimethylated N-terminal peptides that were not identified by HYTANE (Figure 1E). A similar trend albeit with a reduced magnitude was observed for peptides with acetylated N-termini (Figure 1E). Similar results were obtained when LATE and HYTANE were applied to the *E. coli* N-terminome (Figure S3). In these two experiments LATE improved the overall coverage of the N-terminome, but the number of identified peptides with LATE was slightly lower than the number of peptides identified by HYTANE. We noticed that this reduction was more pronounced for acetylated N-termini compared to dimethylated ones, as this may be due to the stronger hindrance of N-terminal amine ionization by the covalently bound acetyl group (Cho *et al*, 2016). We concluded that LATE is better suited for studying proteolysis rather than the natural acetylation of N-terminome, yet it provides valuable information and unique identifications on both.

Next, we applied LATE to study caspase-3 proteolytic activity *in vitro*. The intracellular proteome of HCT116 cells was extracted under native conditions and incubated with recombinant caspase-3 or buffer control at 37°C. Equal aliquots were sampled at 1, 6, or 18 hours of incubation and were subjected to N-terminal enrichment using LATE and HYTANE (Figure 2A). Caspase-3-treated samples and control samples were labeled with heavy and light dimethylation, respectively. Successful enrichment of the N-terminome was obtained with both LATE and HYTANE (Figure S4). Peptides with a high relative abundance ratio (caspase-3/control) are putative products of caspase-3 cleavage. Yet, since caspase-3 might activate other proteases (including other caspases), not all putative cleavages are necessarily caspase-3 cleavages. The distribution of the caspase-3/control abundance ratio in the log_2_ scale was distributed around zero (i.e., no change), with the frequency of peptides with a high caspase-3/control abundance ratio increasing overtime in both methods (Figure 2B). To distinguish the background proteolysis from the caspase-3 cleaved proteins, we considered only peptides that had a caspase-3/control ratio higher than 2 (Log_2_ of 1) as an indication for putative caspase-3 cleavages. Following this, we identified over 900 different cleavage sites after 18 hours of incubation in each one of the methods with more than 200 sites identified at all time points (Figure 2C, and Tables S2 and S3). Caspase-3-mediated proteolysis has been studied extensively and its specificity is well defined (Walsh *et al*, 2008; Zhan *et al*, 2002, 3; Lüthi & Martin, 2007). The sequence logo of the 8 amino acids (P4-P4’) flanking the reported caspase-3 cleavages in the MEROPS peptidase database (Rawlings *et al*, 2012) shows a very similar motif to the logos obtained from our data (Figure 2E). By including only cleavages with a caspase-3/control ratio of ≥2 that were identified only in the caspase-3 treated samples and occurred after Asp or Glu, we selected the most credible caspase-3 cleavage sites. This way, we identified 721 additional putative caspase-3 cleavage sites with LATE that could not be identified with HYTANE (Figure 2D). By combining the results of both methods, we achieved a ∼50% increase in the total number of putative cleavages obtained by HYATNE alone, reaching a total of 2154 putative cleavages enriched in the caspase-3 treated samples.

**Figure 2.**
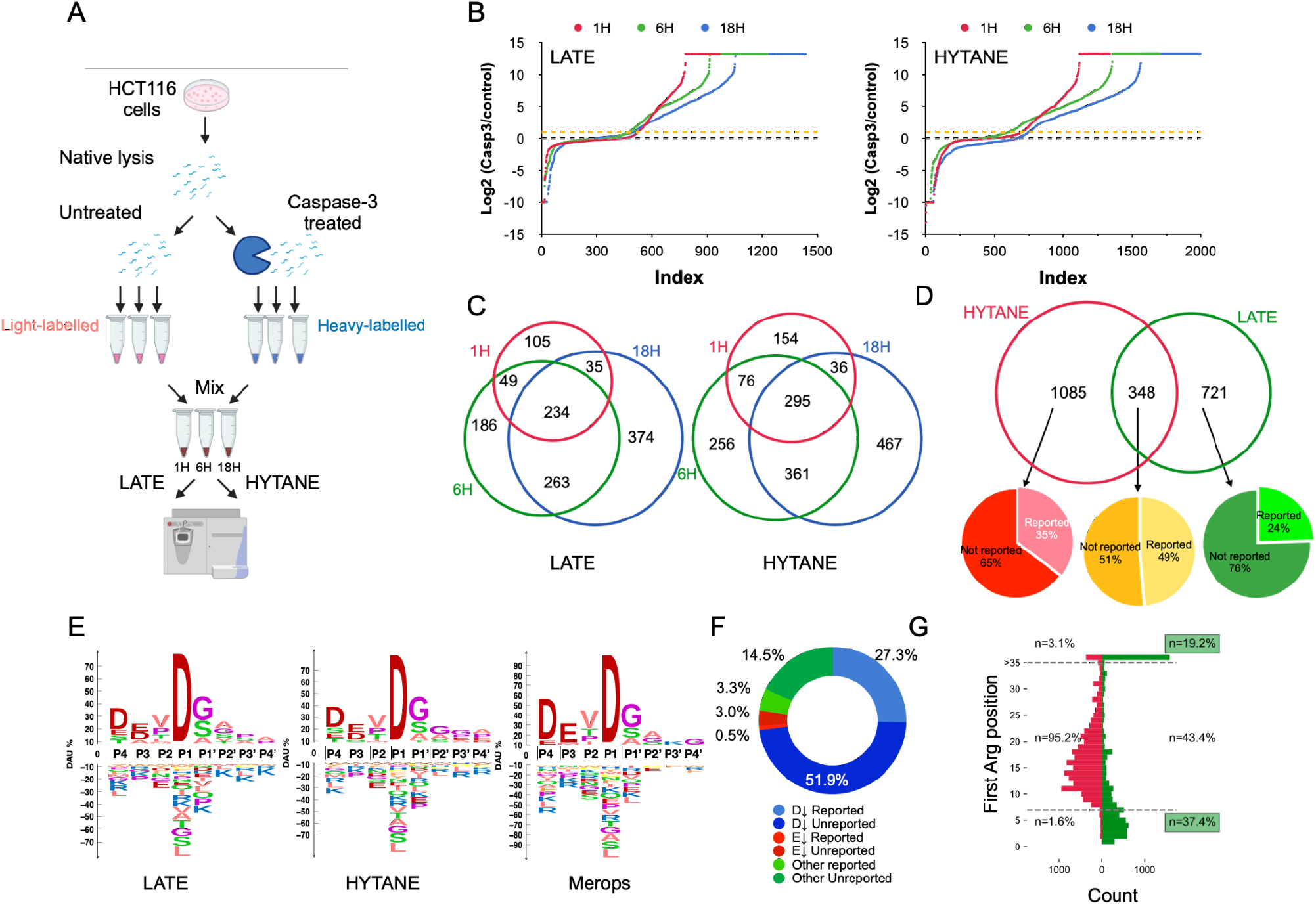
N-terminomics of caspase-3 cleavages *in vitro*. **A**. *In vitro* experimental scheme. **B**. Ranked ratios of dimethylated peptides showing the accumulation trends of peptides with high Log_2_ (caspase-3/control) abundance ratio (>1 as marked by the yellow dotted line) in LATE and HYTANE **C**. Venn diagram of peptides with Log_2_ (caspase-3/control)>1 at each time point in LATE and in HYTANE **D**. Venn diagram of the unique cleavage sites after D or E that were identified only in the caspase-3-treated samples with a caspase-3/control ratio ≥2 in at least one time point. “Reported/not reported” is based on a comparison to published data **E**. Sequence logo of all putative cleavage sites that were identified only in the caspase-3-treated samples with a caspase-3/control ratio ≥2, in comparison to all caspase-3 cleavage found in MEROPS. **F**. Distribution of all putative cleavage sites were identified only in the caspase-3-treated samples with a caspase-3/control ratio ≥2 based on the P1 position of their cleavage motif. **G**. The residue distance distribution of the nearest arginine residue to the identified peptides sequence in LATE (green) and HYTANE (red) experiments.

Of these putative cleavages, 733 (35%) matched previously reported caspase-3 cleavages in TopFind (Fortelny *et al*, 2015), CASBAH (Lüthi & Martin, 2007), DegraBase (Crawford *et al*, 2013), MEROPS (Rawlings *et al*, 2012) and a recent caspase-3 N-terminomics study (Araya *et al*, 2021), while 1421 (65%) have not been reported so far. Some of these unreported cleavages occurred in 372 proteins that were not known previously as caspase-3 substrates (Tables S2 and S3). Interestingly, the proportion of unreported/novel putative caspase-3 cleavages obtained with LATE was higher than with HYTANE (74% vs 63%, respectively) (Figure 2D, Table S2 and Table S3). We were able to identify 91 additional putative caspase-3 cleavage sites with Glu at P1 (Tables S2 and S3), consistent with caspase-3 ability to cleave after Glu residues (Seaman *et al*, 2016). To evaluate the additional contribution of LATE to the results that can be achieved by ArgC-based N-terminomics methods like HYTANE, we compared the sequences of 7 to 35 amino acids downstream to the identified cleavage sites by each method. HYTANE requires an arginine at one of these 7-35 amino acid segments while LATE requires a lysine at these positions. As shown in Figure 2G, more than 50% of LATE identifications consisted of sequences containing Arg residues outside positions 7 to 35 downstream to the cleavage site and a similar trend was found in the location of Lys residues in the peptide identified by each method (Figure S5). This demonstrates LATE’s ability to identify cleavage sites that cannot be found with HYTANE and vice versa.

Next, we used the same combined N-terminomics approach to study caspase-3 substrates in cells. To this end, HCT116 cells were transfected with a bicistronic caspase-3/GFP plasmid and FACS-sorted to generate caspase-3 overexpressing cells (GFP-positive) and non-transfected cells (no GFP). Caspase-3 activity was induced by ABT-199 in both populations (Figure 3A). Overexpression of caspase-3 accelerated apoptosis in HCT116 compared to non-transfected cells in response to ABT-199 (Figure S6B). We used LATE and HYTANE to characterize the ORF- and neo-N-terminal peptides in those cells. This way, changes in proteolytic processing between these cell populations could be directly attributed to caspase-3. The majority of identified peptides were ORF N-terminal peptides, and most of those were acetylated (Figure 3B) with ∼57% of the total identifications in HYTANE and ∼43% in LATE. Yet, the number of neo-N-terminal peptides identified was similar in both methods (Figure 3B) and the sequence logos of those peptides were similar to each other with strong dominance of Asp at the P1 position (Figure 3C). The sequence logos were very similar to those we obtained for caspase-3 *in vitro* and the caspase-3 sequence logo based on MEROPS (Figure 2E). This indicates the dominance of caspase-3 processing among the neo-N-terminal peptides. Excluding the N-terminal peptides of ORFs or alternative translation initiation sites and N-terminal peptides resulting from known protein processing events, such as signal peptide or prodomain removals, we detected 2330 neo-cleavage sites for 1117 different proteins. Although with LATE fewer PSMs were obtained than with HYTANE (Figure 3B), it enabled the identification of more neo-N-terminal peptides (1407) than HYTANE (1127) (Figure 3D and Tables S4 and S5).

**Figure 3.**
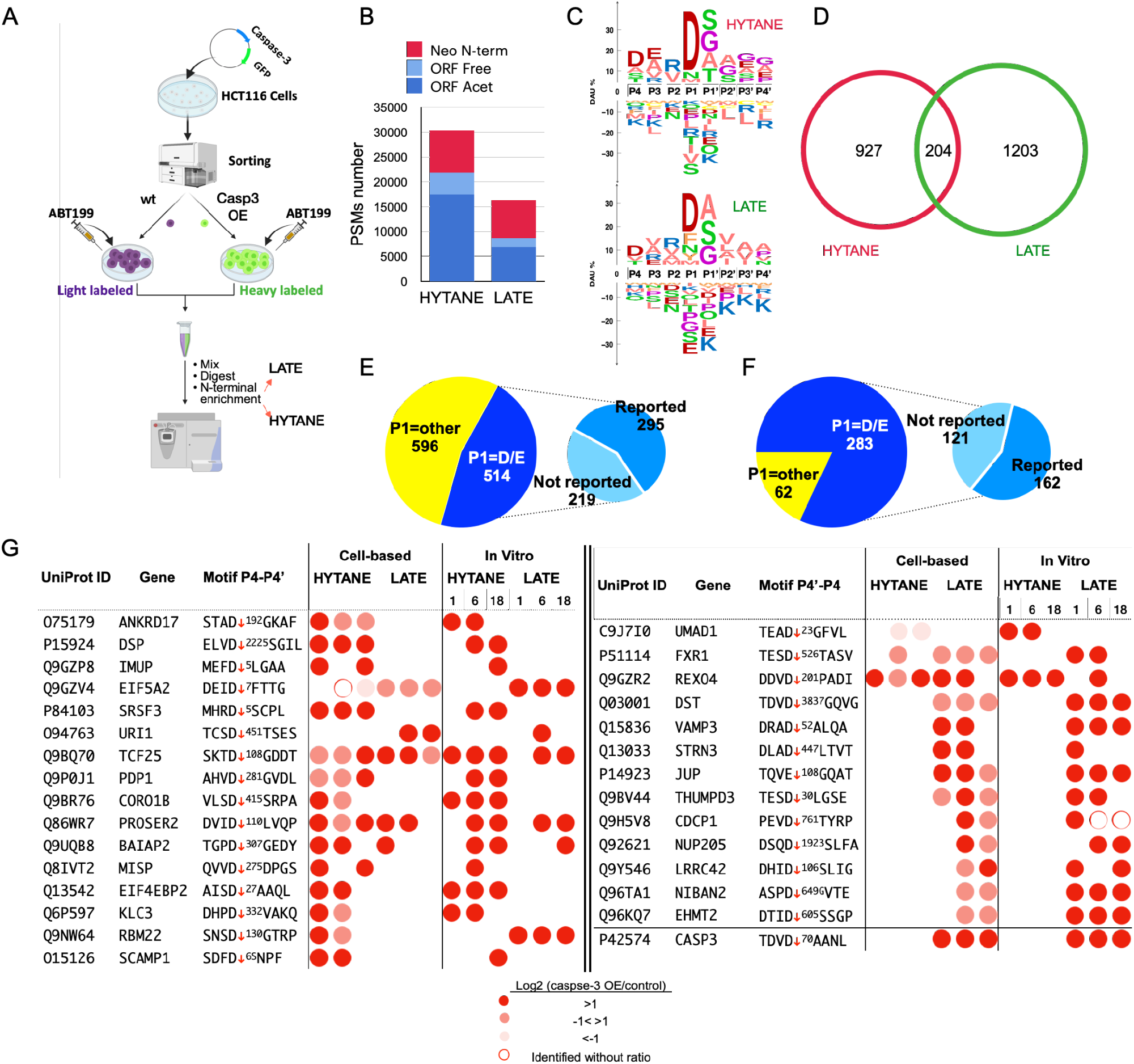
N-terminomics of caspase-3 cleavages in cells. A. Cell-based experimental scheme to study caspase-3 mediated cleavages in HCT116 following treatment with ABT-199 **B**. Number of N-terminal peptides identifications following HYTANE and LATE enrichments **C**. Sequence Logo of neo-N-terminal peptides cleavage motifs by HYTANE (top) and LATE (bottom) **D**. Venn diagram of the combined number of neo cleavage motifs identified by HYTANE (red) and LATE (green). **E**. The number of putative caspase cleavage motifs (with P1=D or E) out of the total neo-N-terminal cleavage motifs that were identified. The categorization of reported/not reported is based on a comparison to published data (TopFind etc). **F**. The number of putative caspase cleavage motifs (with P1=D or E) that had a high caspase-3/control abundance ratio (Log_2_≥1) out of the total neo-N-terminal cleavages identified **G**. Novel caspase-3 substrates and their cleavage sites. The cleavage site (caspase-3/control) abundance ratio is indicated by color.

To chart the changes in the N-terminome we combined the identifications obtained by both methods and considered only those that were identified at least twice. Altogether, 1127 neo-cleavage sites were identified and from these, 519 (∼46%) were cleavages after Asp or Glu corresponding to the caspase family specificity (Figure 3E). Of these putative caspase cleavages, 221 (∼42%) were at sites that were not reported so far in proteomic studies of caspases (Figure 3E and Tables S4 and S5). Some of these unreported cleavages occurred in 65 proteins that were not known so far as caspase-3 substrates (Table S6). A closer look at the Log_2_ (caspase-3/control) abundance ratios of the 298 caspase cleavage sites previously reported in other proteomics studies shows that the majority are larger than 1, and a significant number of the Log_2_ ratios are within the range of - 1 to 1 (Figure S8). Caspase-3 was activated by ABT-199 (Souers *et al*, 2013) in both the caspase-3 over-expressing and control cells, hence there is expected to be no or only a small difference between the two treatments for substrates that are cleaved effectively by the endogenous caspase-3. Therefore, in this work, we also report cleavages without large fold change as putative caspase-3 cleavages. Of note, despite the difference in background proteolysis between the cell-based (Figure 3A) and the *in vitro* experiments, when we compared the abundance ratios of the same substrate cleavage across the two experiments, they showed similar trends (Figure S8). Moreover, for substrate winnowing of caspase-3-related cleavages, we filtered the data further and considered only cleavages that show at least a 2-fold increase in the cells that overexpressed caspase-3 (Figure 3F). We identified several putative caspase-3 cleavages that have not been reported previously, either in the various studies related to apoptosis or in caspase-3-related proteomics studies (Lüthi & Martin, 2007; Fortelny *et al*, 2015; Crawford *et al*, 2013; Araya *et al*, 2021) (Figure 3G and 3F and Table S6). Of the 345 cleavages that show significant change, 283 (over 80%) were with caspase specificity. Of these, 162 (57%) were at sites that were previously reported. and 121 (43%) are at sites that were not reported. From these unreported 121 sites we selected several substrates and demonstrated that they are cleaved by caspase-3 *in vitro* (Figure 3G and Table S2 and S3). All of the novel substrates and their cleavage sites are shown in Table S6. Of note, one of the unreported cleavage sites identified in all of our experiments, exclusively by LATE, is a cleavage site at position 70 of caspase-3 itself (Figure 3G).

We used gene ontology (GO) enrichment analysis to compare the list of unreported putative caspase-3 substrates that we identified to previously reported substrates from apoptosis and caspase-related proteomic studies. The new putative substrates show a pattern highly similar to the proteins previously reported to be cleaved caspase-3, thus demonstrating that the new cleavages come from substrates with similar biological function to the already known ones (Figure S9). The number of potential caspase-3 cleavages and substrates identified in the cell-based experiments was smaller than that obtained in the *in vitro* experiments. In order to understand whether there is a structural basis for the differences between the cleavages obtained in the cell-based and *in vitro* experiments, we compared the secondary structure contexts and solvent accessibility around the cleavage sites. This comparison revealed very similar secondary structure contexts for the two experiments (Figure S10A) but a significant difference in the relative solvent accessibility (Figure S10B), with the cleavages identified in cells occurring in more exposed regions than those obtained *in vitro*. This might reflect the impact of cell lysis in the *in vitro* experiments that alters some protein structures and disrupts cellular compartmentalization that is kept under control in the cell-based experiments.

One of the advantages of the negative selection N-terminomics methods is that in addition to the enrichment of the neo-N-terminal peptides, these methods also allow the characterization of the ORF N-terminal peptides (Figures 1E and 1F) (Kleifeld *et al*, 2010). We used the same approach applied to the neo-N-termini peptides and combined the ORF N-terminal peptides identification obtained by LATE and HYTANE. In our search we included these known modifications of protein N-termini: removal of the initiator methionine (Giglione *et al*, 2015, 2004) and Nt-acetylation (Ree *et al*, 2018). Together, the ORF N-terminal peptides of 2349 proteins were identified (Table S7). Similarly to published data (Arnesen *et al*, 2009), 74% of the peptides were Nt-acetylated, 26% with free N-termini (Nt-free) and 70% of all peptides underwent methionine excision regardless of their Nt-acetylation status (Figure 4A). Additionally, the frequencies of the first and second amino acids in Nt-acetylated and Nt-free ORF peptides (Figure S11) were similar to those found in other proteomic studies (Arnesen *et al*, 2009).

**Figure 4.**
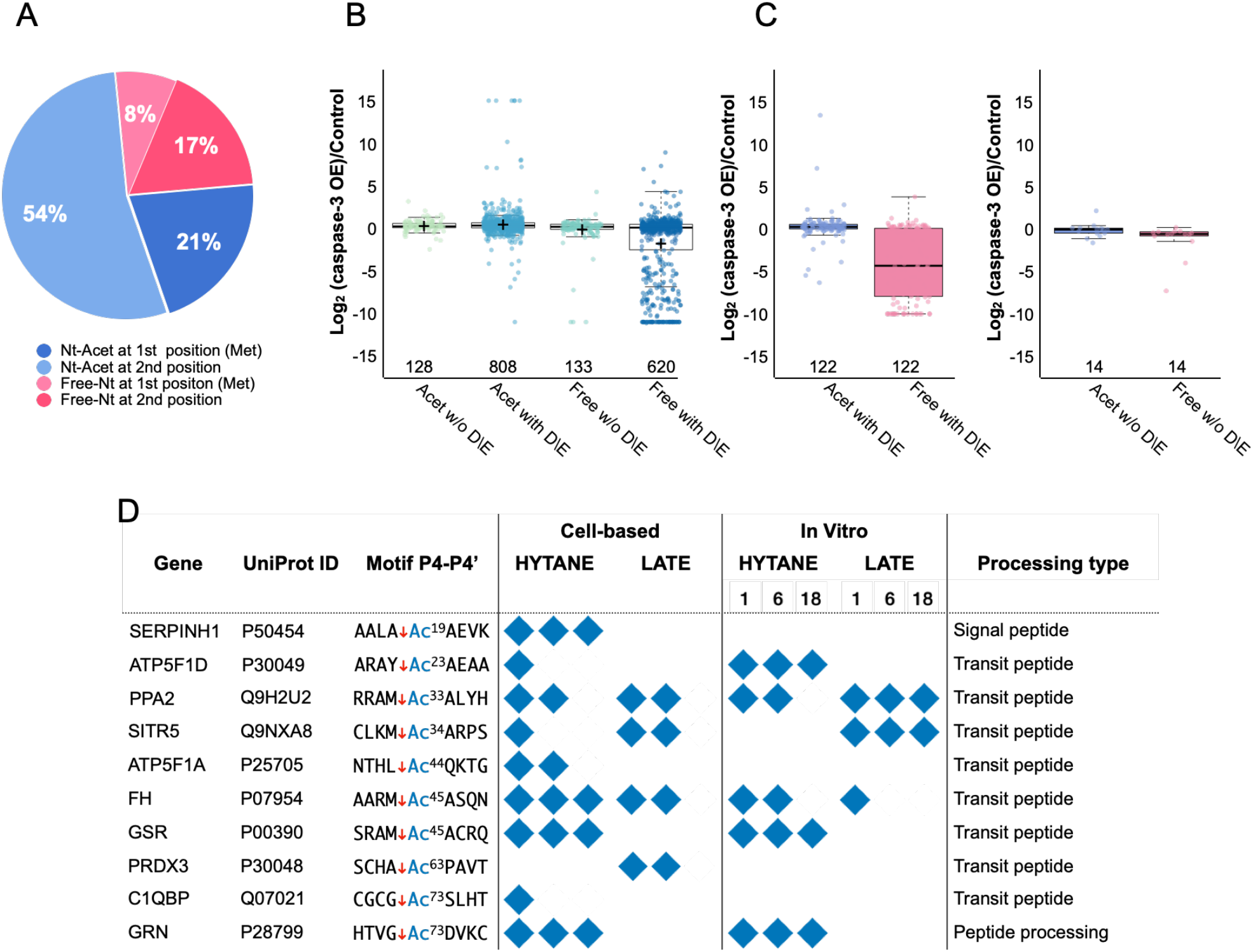
Acetylation of both ORF and neo N-terminal peptides. **A**. The ORF N-terminal peptides identified in HCT116 cells overexpressing caspase-3 and their control were categorized based on the presence of initiation methionine (marked as 1st Met) or its removal (marked as 2nd position), the presence of Nt-acetylation (“Nt-Acet”; in shades of blue) or its absence (“Free”; in shades of pink). **B**. The abundance ratio distribution of the ORF N-terminal peptides in cells overexpressing caspase-3 and their control, based on their N-terminal state (free/acetylated) and the presence of Asp or Glu residue in their sequence. Center lines show the medians; box limits indicate the 25th and 75th percentiles; whiskers extend 1.5 times the interquartile range from the 25th and 75th percentiles; crosses represent sample means; data points are plotted as open circles. n = 128, 808, 133, 620 sample points **C**. The abundance ratio distribution of the 122 ORF N-terminal peptides that were identified both as Nt-acetylated and with free N-terminal and contain Asp or Glu in their sequence. Center lines show the medians; box limits indicate the 25th and 75th percentiles; whiskers extend 1.5 times the interquartile range from the 25th and 75th percentiles; crosses represent sample means; data points are plotted as open circles. **D**. Post-translational neo-Nt-acetylation sites that were identified at N-termini generated following known proteolytic processing of precursor proteins as indicated in Uniprot. The blue diamonds represent identification in biological repeats or time points.

Next, we checked if the nature of the ORF N-terminal modification affects the susceptibility of ORF N-termini to cleavage by caspase-3. As can be seen in Figure 4B and Figure S12, the abundance ratio of Nt-acetylated ORF peptides was distributed around zero regardless of the presence of Asp or Glu in their sequence. A similar trend was seen for Nt-free ORF peptides that do not contain potential caspase-3 cleavage sites. However, Nt-free ORF peptides with Asp or Glu in their sequence were more susceptible to cleavage by caspase-3 and many of their ratios deviated from a normal distribution and demonstrated lower abundance ratios in caspase-3 overexpressing cells relative to the control cell. Among these ORF N-terminal peptides, 136 were identified and quantified both in the Nt-acetylated form and in the free N-terminal form. Of these peptides, 122 contained Asp or Glu residues, thus containing putative caspase cleavage motifs. The abundance ratios of these peptides in their N-terminal acetylated form were spread around zero while the abundances of their N-terminal free forms were much higher in the control (Figure 4C). The abundance ratio of the 14 peptides that did not contain Asp or Glu in their sequence was distributed around zero regardless of the nature of their N-terminus (Figure S13). This supports the notion that Nt-acetylation may have a protective and stabilizing role as has been suggested in some studies (Mueller *et al*, 2021). Nt-acetylation is defined mostly as a co-translational modification, but there are a few examples of post-translational Nt-acetylation (Aksnes *et al*, 2015; Drazic *et al*, 2018, 80; Helsens *et al*, 2011; Helbig *et al*, 2010; Linster & Wirtz, 2018). To investigate the presence of post-translational Nt-acetylation that is not at the protein’s ORF N-terminus, we checked the N-terminal peptides that were generated following known proteolytic processing of proteins. These processing events include signal/transit peptide or propeptide removal and peptide or precursor protein chain cleavages that are part of protein maturation or activation. By combining LATE and HYTANE results, we identified 329 post-translational proteolytic processing sites reported in UniProt for 262 proteins (Table S8). Of these, 11 neo-Nt-peptides from 11 proteins were found with Nt-acetylation (Figure 4C) as well as with free N-terminus. The neo-acetylated-Nt-peptides identified were mainly from the mitochondrial proteins and derived from the removal of the mitochondrial transit peptide. This type of proteolytic processing is done after the protein is imported into the mitochondria and reaches the mitochondrial matrix (Friedl *et al*, 2020) or inner membrane (Ieva *et al*, 2013). Next, we looked for additional hints of post-translational neo-Nt-acetylation that occur following proteolytic cleavages not annotated in UniProt (The UniProt Consortium, 2021) and not reported as an alternative initiation site in TopFind (Fortelny *et al*, 2015).

We found 371 such neo-Nt-acetylation motifs in 329 different proteins (Table S9). Of these neo-Nt-acetylation sites, 111 were after Asp and fit caspase-3 specificity. Interestingly, 75 sites out of these 111 putative sites have previously been reported as apoptosis-related and include several known caspase-3 cleavages of translation-related proteins such as EIF5A, EIF3J and EIF4H. To further check the link between neo-Nt-acetylation sites in the list and caspase-3 cleavage, we used the winnowing workflow shown in Figure 5A.

**Figure 5.**
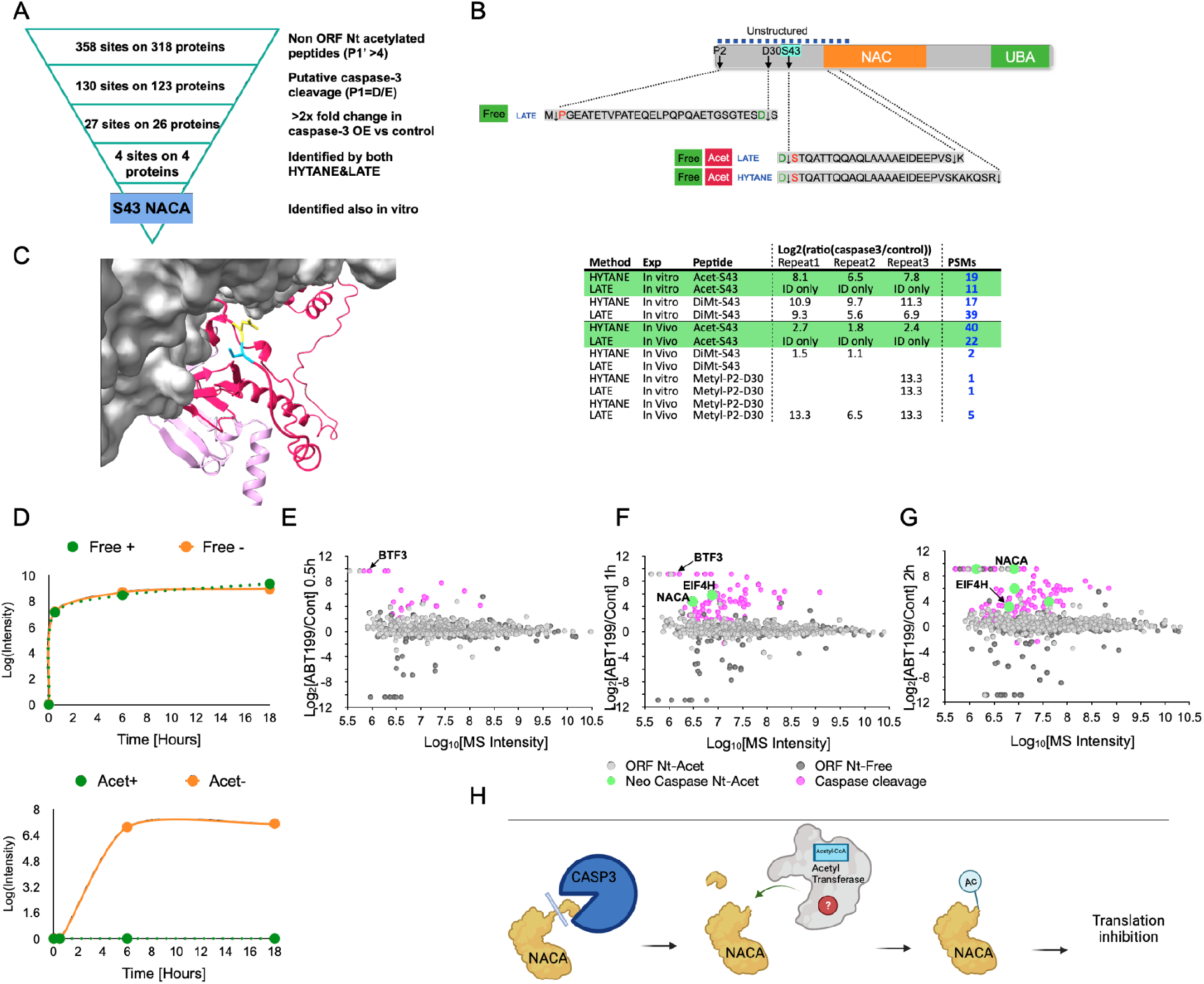
Neo-Nt-acetylation of NACA of ORF and neo-N-terminal peptides. A. Winnowing of acetylated neo-N-terminal peptides identified in HCT116 cells. **B**. The N-terminal peptides of Nascent polypeptide-associated complex subunit alpha (NACA) identified in HCT116 cells. Cleaved peptides sequences and modifications are shown on the top and the PSMs numbers in the different experiments are on the bottom **C**. The position of NACA cleavage (in yellow and light blue) by caspase-3, before Ser43, lead to the formation of a neo-Nt-acetylated form of NACA. The structure model is based on the structure of the ribosome-nascent chain containing an ER signal sequence in a complex with the NAC complex (PDB:7QWR). The ribosome and RNA are in grey. The full NACA (in dark pink) and BTF3 (NACB) (in light pink) structures are based on AlphaFold (Varadi *et al*, 2022) prediction and were aligned on top of the original partial structures. **D**. Time-course of *in vitro* NACA cleavage and Nt-acetylation. The upper panel shows NACA cleavage at Ser43 by caspase-3 with (green) or without (orange) lysate dialysis before caspase-3 addition. The lower panel shows NACA neo-acetylation at Ser43 after caspase-3 cleavage with (green) or without (orange) lysate dialysis before caspase-3 addition **E-F**. Time-course N-terminomics of HCT-116 cells treated with ABT-199 or DMSO (c20ontrol). Log_2_ abundance ratios for individual peptides are plotted on the y-axis and the corresponding total peptide MS intensities are shown on the x-axis. The circle colors correspond to the peptide type: ORF Nt-Acetylated (light gray), ORF peptides with Nt-free (dark gray), caspase-generated neo-Nt peptides (pink), and neo-Nt-acetylated peptides (light green). **H**. The suggested order of events for NACA neo-Nt-acetylation following caspase-3 processing.

Consequently, we focused on the proteolytic cleavage and neo-Nt-acetylation of nascent-polypeptide-associated complex A (NACA). The MS/MS identifications of this neo-Nt-acetylated peptide by LATE and HYTANE had high scores, high peptide sequence coverage and provided conclusive evidence for the presence of Nt-acetylation on Ser43 (Figure S13). NACA is a subunit of the hetero-dimeric ribosome-associated (Rospert *et al*, 2002) nascent-polypeptide-associated complex (NAC). The NAC complex is highly conserved among eukaryotes, and when associated with the ribosome it interacts with nascent polypeptide chains and mediates their sorting to different cellular compartments (Jomaa *et al*, 2022). In addition to the neo-Nt-acetylation of the peptide starting at Ser43, we found several neo-N-terminal peptides of NACA (Figure 5B upper part). The majority of these cleavages were also reported in other apoptosis-related studies (Lüthi & Martin, 2007; Fortelny *et al*, 2015; Crawford *et al*, 2013).Interestingly, in the cell-based experiments, this peptide was identified almost exclusively in its acetylated form with a total of 62 PSMs vs 2 PSMs for the free form, while in the *in vitro* experiments the total PSM numbers of both forms were similar. The location of this cleavage is within the N-terminal unstructured region of NACA which makes it suitable for caspase cleavage. Based on a recent structure of the NAC complex bound to the ribosome (Jomaa *et al*, 2022) this region remains accessible even when NACA is in complex with BTF3 (nascent-polypeptide-associated complex B - NACB) and the ribosome (Figure 5C). The NACA Nt-acetylated Ser43 peptide was identified *in vitro* only after the addition of active caspase-3, which allowed us to study its generation process. We used HCT116 cell lysate and half of it was subjected to dialysis to reduce acetyl-CoA concentration and prevent the acetylation reaction. Following the addition of active caspase-3, aliquots from both dialyzed and non-dialyzed lysates were sampled at different time points and subjected to N-terminal enrichment and MS analysis. As shown in Figure 5D, the dialysis did not affect caspase-3 cleavage and the appearance of NACA neo-Nt-Ser43. This peptide was already identified 30 minutes after caspase-3 addition and was generated at an identical rate and intensity in both samples. The NACA neo-Nt-acetylated Ser43 peptide appeared at a later time point (6 hours) but only in the non-dialyzed sample (Figure 5E). Next, we checked if and at which stage NACA neo-Nt-acetylation can be observed during apoptosis. HCT116 were treated with ABT-199 and DMSO (control). Cells were collected from the ABT-treated and control cells after 0.5, 1, and 2 hours after addition of ABT-199 or DMSO and were then subjected to N-terminal peptide enrichment using HYTANE. No cell death was observed until 2 hours of treatment (Figure S6B). As shown in Figures 5E-G, NACA neo-Nt-acetylation at Ser43 was already observed after 1 hour of treatment (marked in green). Besides NACA, neo-Nt-acetylation that precedes cell death was also identified on the translation elongation factor eIF4H after it was cleaved by caspase (at Ser94). Interestingly, we also observed a cleavage (occurred after 0.5 hours of treatment) of BTF3, the protein that forms the ribosome-adjacent NAC with NACA (Gamerdinger *et al*, 2019; Jomaa *et al*, 2022). Together with the rest of our results, this, for the first time, reveals the presence of post-translation Nt-acetylation that is directly linked to caspase-3 proteolytic processing (Figure 5H).

## Discussion

The use of multiple proteolytic enzymes in bottom-up proteomics studies is known to improve the number of protein identification and sequence coverage (Swaney *et al*, 2010; Giansanti *et al*, 2016). Similar improvements were also shown in several terminomics studies that combined trypsin with other proteases while using the same terminal peptides enrichment methodology (Bell *et al*, 2019; Zhang *et al*, 2018). Here we describe LATE, a simple methodology that is based on digestion with LysN for N-terminome characterization. LysN digestion has been used for the identification of ORF Nt-acetylated peptides (Du *et al*, 2020; Zhang *et al*, 2009) but not for free ORF N-terminal peptides nor neo-N-terminal peptides generated by proteolysis. A key element of LATE is the N-terminal-specific labeling by dimethylation, which also allows the use of isotopic labels and quantitative analysis. Unlike the few other reported N-terminal specific labeling reagents, like TMPP (Bertaccini *et al*, 2013) or 2-pyridinecarboxyaldehydes (Griswold *et al*, 2019; MacDonald *et al*, 2015), the required reagents are inexpensive and the labeling efficiency is very high and consistent. We demonstrated the usefulness of LATE combined with HYTANE, in the investigation of caspase-3 mediated cleavages during apoptosis. This subject has been extensively studied, particularly by terminomics methodologies (Mahrus *et al*, 2008; Seaman *et al*, 2016; Demon *et al*, 2009; Agard *et al*, 2012; Julien *et al*, 2016). We identified many novel caspase-3 cleavage sites and substrates as well as some known substrates whose cleavage by caspase-3 had not been previously identified by proteomic analysis. Such known substrates are desmoplakin (DSP) and plakoglobin (JUP) which have been known as caspase-3 targets for over two decades (Weiske *et al*, 2001; Brancolini *et al*, 1998) but their cleavage sites had not yet been identified by proteomics. We identified DSP and JUP cleavage sites as part of a large-scale study for the first time. The novel substrate list includes several proteins that are understudied in the context of apoptosis. For example, the unconventional prefoldin RPB5 interactor 1 (URI1) that is important to DNA stability (Parusel *et al*, 2006) and affects apoptosis upon its phosphorylation (Djouder *et al*, 2007), and Transcription factor 25 (TCF25 or NULP1) which binds the apoptosis inhibitor XIAP (Steen & Lindholm, 2008) and the overexpression of which induces cell death. Our combined system-wide *in vitro* and cell-based experiments allowed us to validate these proteins as caspase-3 substrates. In addition, this thorough N-terminomics characterization enabled us to monitor not just proteolytic cleavage but also changes in protein ORF N-termini. In this respect, our results indicate that Nt-acetylation protects proteins’ N-terminal regions from proteolytic cleavage by caspase-3 during apoptosis. In mammalian proteins, Nt-acetylation can be seen in more than 70% of ORF N-termini, yet its physiological functions are not entirely understood. Nt-acetylation affects the charge of proteins’ N-termini hence it would be expected to modify their function. Initially, it was suggested that a general function for Nt-acetylation is the protection of proteins from proteolytic degradation (Hershko *et al*, 1984; Jörnvall, 1975). Later on, it was suggested that under certain conditions, Nt-acetylation might also be a degradation signal (degron) that marks the protein for ubiquitinylation and its destruction by the proteasome (Hwang *et al*, 2010). We showed that in the context of apoptosis, the presence of Nt-acetylation has a protective effect and prevents cleavage around the protein N-terminal by caspase-3.

Most Nt-acetylation events are thought to be co-translational, but several post-translational Nt-acetylation events were reported too. Among them are the Nt-acetylation of secreted proteins in Apicomplexa (Nyonda *et al*, 2022), cleaved proteins in yeast by an unknown protease (Helsens *et al*, 2011; Helbig *et al*, 2010), transmembrane proteins in the Golgi (Aksnes *et al*, 2015) and cleaved actin (Drazic *et al*, 2018). Nt-acetylation of nuclear-coded proteins imported to the mitochondria has been described but only co-translational ORF Nt-acetylations were identified (Vaca Jacome *et al*, 2015; Bienvenut *et al*, 2012). Here we uncover the Nt-acetylation of several mitochondrial proteins that occur after their transit peptide removal in the mitochondrial matrix. Nt-acetylation of imported nuclear-encoded proteins after the cleavage of their transit peptides was identified in chloroplasts (Zybailov *et al*, 2008; Huesgen *et al*, 2013; Bienvenut *et al*, 2012). Both organelles are thought to have a prokaryotic origin, yet Nt-acetylation is rare in bacteria (Bienvenut *et al*, 2012). This raises the question of which N-acetyltransferase catalyzes the acetylation of mitochondrial proteins. Recently a family of dual lysine and N-terminal acetyltransferases was identified and several of its members were shown to reside within plastids (Bienvenut *et al*, 2020). Mammalian mitochondrial N-acetyltransferase has not been not identified.

We also revealed several caspase-3-generated neo-N-termini that undergo post-translational Nt-acetylation. We focused on and validated the caspase-3-dependence of NACA neo-Nt-acetylation that was dominant in all of our experiments and analyses. Aside from NACA’s role as part of a ribosome-associated complex, it was shown to function as a transcriptional coactivator regulating bone development and hematopoiesis. Hence, its cleavage and Nt-acetylation might have implications on transcription. As part of a ribosome-associated complex, NACA binds to nascent proteins that lack a signal peptide motif as they emerge from the ribosome, while blocking their interaction with the signal recognition particle and preventing mistranslocation to the endoplasmic reticulum (Jomaa *et al*, 2022). Downregulation of NACA, by hypoxia or RNAi, can lead to the initiation of an ER stress response that is followed by caspase-dependent apoptosis (Hotokezaka *et al*, 2009, 2015). The N-terminal region of NACA cleaved by caspase-3 contains a cluster of negatively charged residues (Figure S14 top). This negatively charged region was shown to downregulate ribosome binding of NAC. (Gamerdinger *et al*, 2019). It was suggested that this negative N-terminal region may interact with the positively charged ribosome-binding motifs of both NACA and its partner BTF3 (also known as NACB). Deletion of the N-terminal region increased NACA binding to the ribosome and caused severe translation inhibition (Gamerdinger *et al*, 2019). Therefore, the apoptosis-induced caspase cleavage of NACA is expected to have a similar inhibitory effect on translation. Based on the Nt-acetylation’s stabilizing effect on protein termini, these observations suggest that neo-Nt-acetylation protects the malfunctioning NACA and thus promotes translation inhibition. Interestingly, we also observed a cleavage of BTF3 (Figure 5E and 5F), that appeared at the earliest time point. Similarly, to the cleavage of NACA at Ser43, BTF3 cleavage after Asp176 led to the removal of a cluster of negatively charged residues (Figure S14 bottom). Thus, it is likely that caspase-3 cleavage of BTF3 also increases NAC binding to the ribosome and likewise inhibits translation. We also observed neo-Nt-acetylation following caspase cleavage of the eukaryotic translation initiation factor 4H (eIF4H) (Figure 5E and 5F and Supplementary Table S10). This protein was shown to enhance the activity of the initiation factor complex eIF4F by stimulating the helicase activity of eIF4A (Pelletier *et al*, 2015). The cleavage of eIF4H by caspase-3 between Asp93 and Ser94 lies directly in the middle of its RNA recognition motif and therefore is expected to abolish its activity and inhibit the RNA priming activity of the whole eIF4F. The neo-Nt-acetylation of eIF4H may shield the cleaved protein from additional proteolytic processing, thus maintaining the inhibition of RNA priming and of translation initiation. We also identified caspase-mediated cleavages of several factors that control translation elongation like eIF5A and eIF5A2, as has also been reported before (Bushell *et al*, 2004). Some of these cleavages were observed shortly after apoptosis induction (Supplementary Table S10). This demonstrates that translation inhibition begins during the early phases of apoptosis and that it is achieved by the combination of ribosome blocking, interference with translation initiation, and elongation blocking.

Together, our data reveal an interplay between caspase-mediated proteolysis and post-translational Nt-acetylation. A similar functional interplay was described before for caspase-mediated proteolysis and phosphorylation (Dix et al., 2012; Seaman et al., 2016). It was shown that caspase cleavage can expose new sites for phosphorylation and that phosphorylation sites can enhance caspase cleavage in their vicinity (Dix *et al*, 2012). We found that caspase-3 cleavage can unveil new sites for post-translational Nt-acetylation and conversely that Nt-acetylation protects proteins from caspase cleavage. It would be interesting to combine N-terminomics using LATE and HYTANE with phosphoproteomics to investigate if there is tripartite crosstalk between caspase-mediated proteolysis, phosphorylation, and Nt-acetylation and if these modifications outcomes are additive or antagonistic to each other.

## Significance

N-terminomics methods are proteomics methods aimed at the characterization of protein N-terminal sequences. These include the mature proteins’ N-terminal sequences as well as N-termini generated following proteolysis. We developed a method to isolate N-terminal peptides that is based on proteome digestion with the lysine-specific protease LysN followed by α-amine specific labeling. Our method, called LATE, enables the identification of N-terminal peptides that can not be identified by other N-terminomics methods. By using LATE along with HYTANE, a common N-terminomics method, we substantially improved N-terminome coverage. Applying this approach to study caspase-3 mediated proteolytic changes during apoptosis revealed many unreported caspase-3 cleavage sites that were also validated *in vitro*. Furthermore, these studies revealed, for the first time, an interplay between caspase-mediated proteolysis and Nt-acetylation. We show that Nt-acetylation protects proteins’ N-termini from cleavage by caspase-3 and on the other hand, caspase-3 cleavage can generate new sites for post-translational Nt-acetylation. We suggest that some of these neo-Nt-acetylation sites have a role in translation inhibition which is an early event in apoptosis progression.

## Acknowledgements

This research was supported by ISF grants 1623/17 and 2167/17. While writing this manuscript, R.H. was supported by the Neubauer Fellowship. We thank the Technion Smuler Proteomic Center and especially Ilana Navon for their ongoing support and help with MS measurements.

## Author contributions

Conceptualization, O.K and R.H; Methodology, O.K and R.H; Investigation, R.H, L.S, D.B, A.R and T.L.; Writing – Original Draft, R.H and O.K; Writing – Review & Editing, R.H, A.R., L.S, T.L and O.K; Supervision – O.K.

