## Supplemental Figures S1-S14 and Table S11 for "In-Depth Characterization of Apoptosis N-terminome Reveals a Link Between Caspase-3 Cleavage and Post-Translational N-terminal Acetylation"

**List of items:**

Figure S1: Human ORF N-terminal peptides that can be identified by different proteases.
Figure S2: Depletion of internal peptides and enrichment of N-terminal peptide by HYTANE of HACAT cells
Figure S3: Comparison of *E. coli* N-terminal peptides identified by HYTANE and LATE (including Free and acetylated)
Figure S4: N-terminal peptides enrichment in the *in vitro* caspase-3 experiments
Figure S5: The residue distribution of the nearest lysine residue to the identified peptides sequence in LATE and HYTANE.
Figure S6: HCT-116 cells viability in response to ABT-199
Figure S7: Ratio distribution of putative caspase-3 cleavages in cell-based experiments
Figure S8: Putative caspase-3 cleavages ratio comparison between *in vitro* and cell-based experiments
Figure S9: GO enrichment analysis of unreported caspase-3 substrates
Figure S10: caspase-3 cleavage motif structure analysis
Figure S11: Nature and frequency of amino acids at positions 1 and 2 in ORF N-terminal peptides
Figure S12: The ratio distribution of ORF-N-terminal peptides as a function of initiation methionine and Nt-acetylation and the presence of Asp or Glu residues
Figure S13: MS/MS of Nt-acetylated NACA Ser43 peptide by LATE and HYTANE
Figure S14: NACA and BTF3 sequences

Table S1: LATE labelling - Provided in attached Excel file
Table S2: In vitro cleavages identified by LATE - Provided in attached Excel file
Table S3: In vitro cleavages identified by HYTANE- Provided in attached Excel file
Table S4: In cells cleavage sites identified by LATE - Provided in attached Excel file
Table S5: In cells cleavage sites identified by HYATNE - Provided in attached Excel file
Table S6: Summary of neo-N-term peptides identified in cells - Provided in attached Excel file
Table S7: Summary table of ORF N-terminal peptide identified in cells - Provided in attached Excel file
Table S8: Known processing sites with free and acetylated N-terminal identified in cells - Provided in attached Excel file
Table S9: Neo-Nt-acetylated peptides - Provided in attached Excel file
Table S10: Time-resolved N-terminomics of HCT116 cell early apoptosis provided in the attached Excel file
Table S11: Search parameters for different MS/MS searches (see in page 15)
Table S12: Results of the GO enrichment analysis used provided in the attached Excel file
Table S13: PSM In vitro LATE-HYTANE provided in the attached Excel file
Table S14: PSM In vivo LATE-HYTANE provided in the attached Excel file
Table S15: PSM Time course ABT199 provided in the attached Excel file

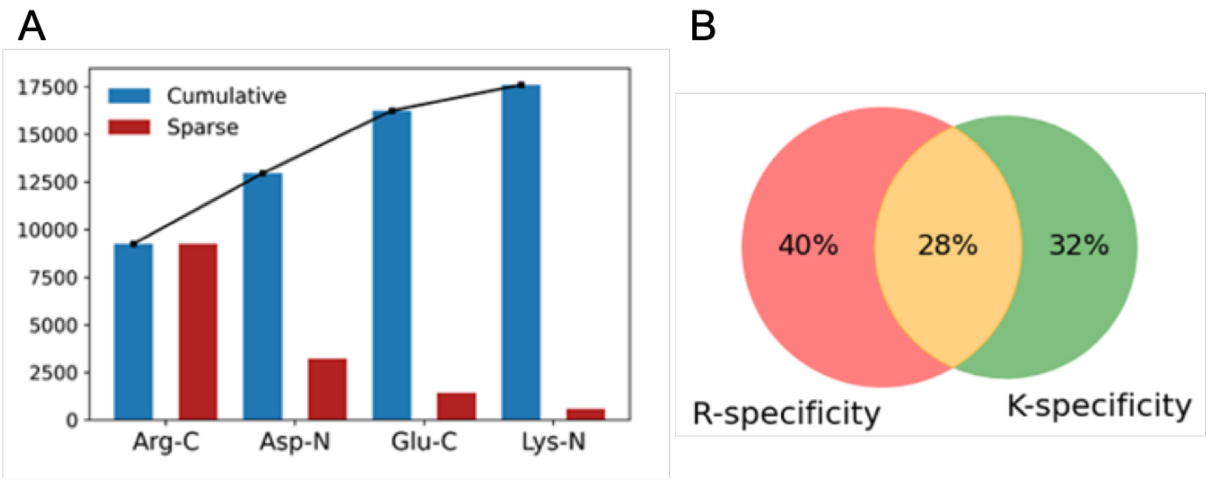


**Figure S1: Human ORF N-terminal peptides that can be identified by different proteases. A**. *In Silico* analysis of the theoretical number of ORF N-terminal peptides of the human proteins (SwissProt, 20198 sequences) that can be identified (peptide length 7-35 amino acid) following digestion with ArgC alone and when ArgC is combined with parallel digestion with other proteases. Blue bars represent the combined number of peptides that can be identified, and red bars represent the specific addition of each protease. **B**. *In silico* analysis of the number of human proteins ORF N-terminal peptides that can be identified following digestion with protease with ArgC-like specificity (red) and LysN (green).

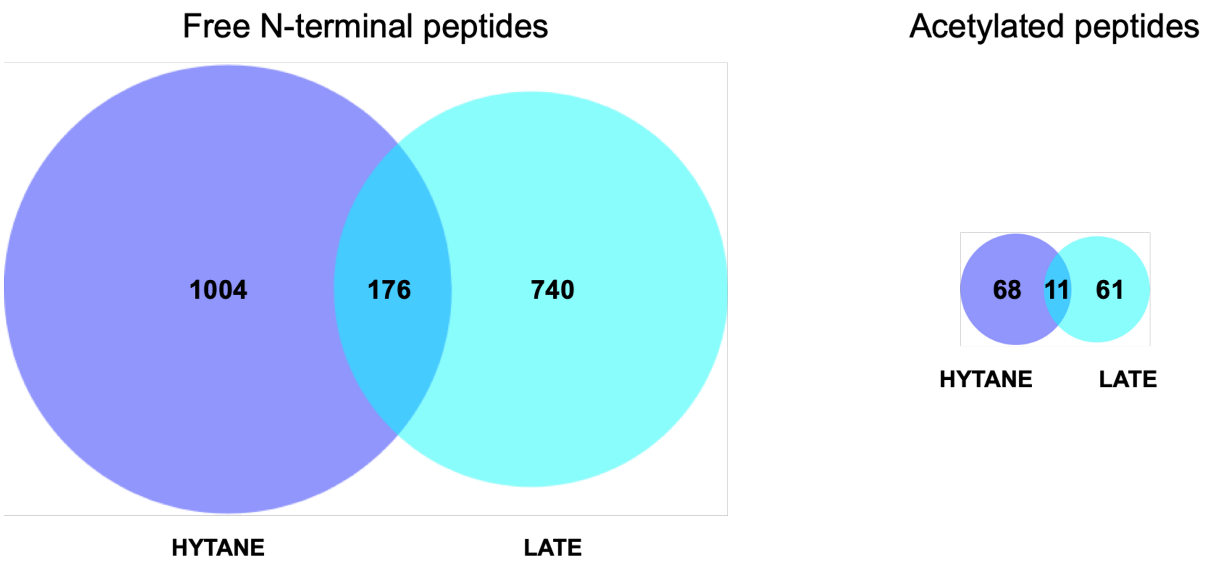


**Figure S2: Comparison of *E. coli* N-terminal peptides identified by HYTANE and LATE.** The number of *E. coli* peptides with free N-terminal (left) or acetylated N-terminal peptides (right) identified by HYTANE (blue) and LATE (cyan),

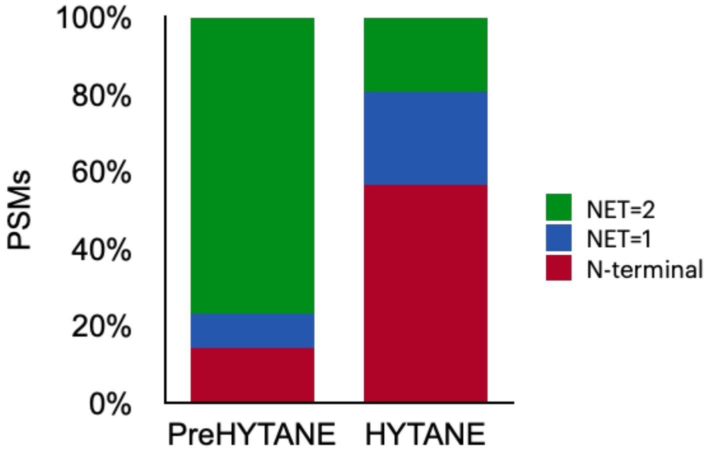


**Figure S3: Depletion of internal peptides and enrichment of N-terminal peptide by HYTANE of HACAT cells.** Peptide-spectrum match (PSM) number of ArgC internal peptides (green) versus the PSM number of ORF (burgundy) and neo N-terminal peptides (blue) before (PreHYTANE) and after HYTANE.

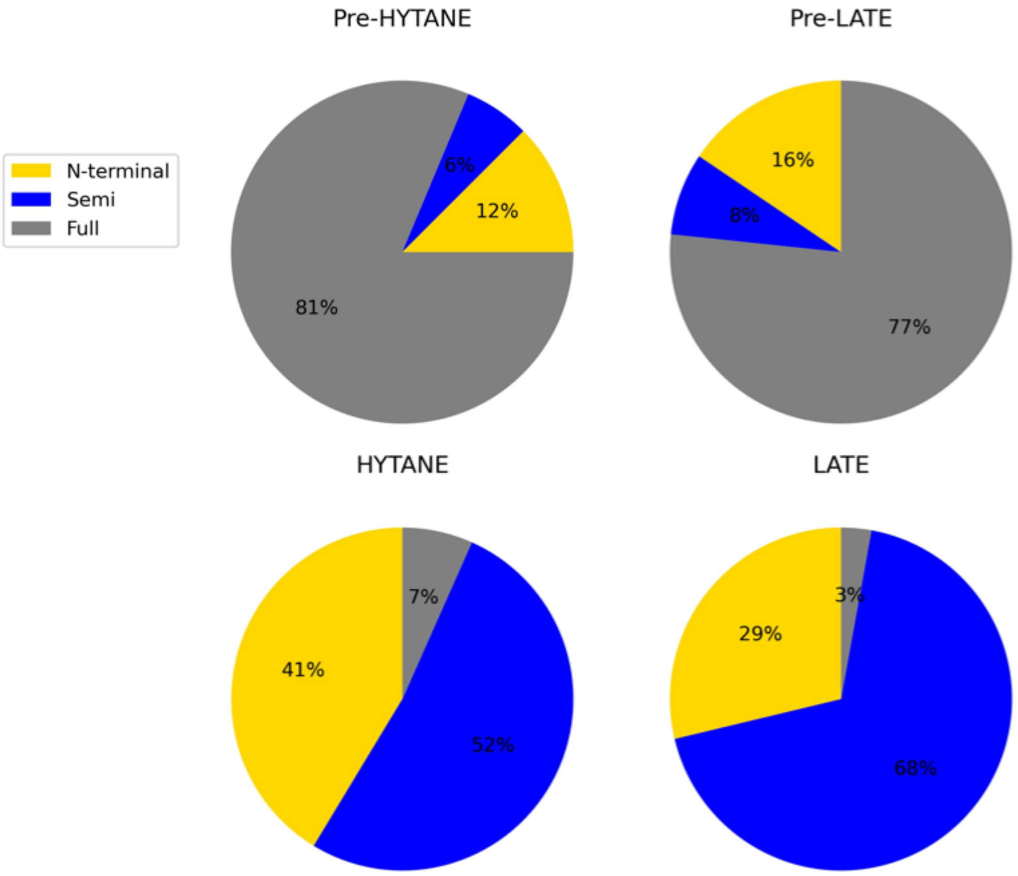


**Figure S4: N-terminal peptides enrichment in the *in vitro* caspase-3 experiments.** The peptides that were identified in all of the *in vitro* caspase-3 experiments were categorized based on their number of enzymatic (NET) and start position. Peptides starting at the ORF position 1 or 2 were defined as ORF N-terminal peptides (yellow). Peptides with NET=2 were defined as "Full" (grey) and those with NET=1 were defined as "Semi" (blue).

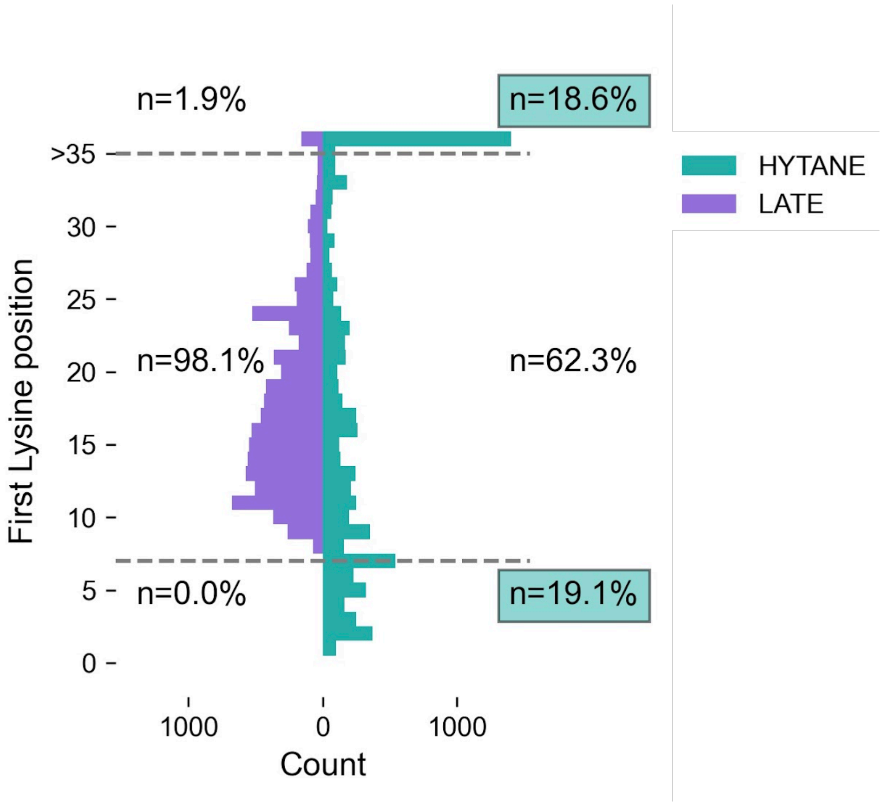


**Figure S5: The residue distribution of the nearest lysine residue to the identified peptides sequence in LATE and HYTANE**. Based on the peptide identifications in the in vitro caspase-3 experiment with LATE (purple) and HYATNE (cyan), the relative distance to the nearest lysine residue was calculated.

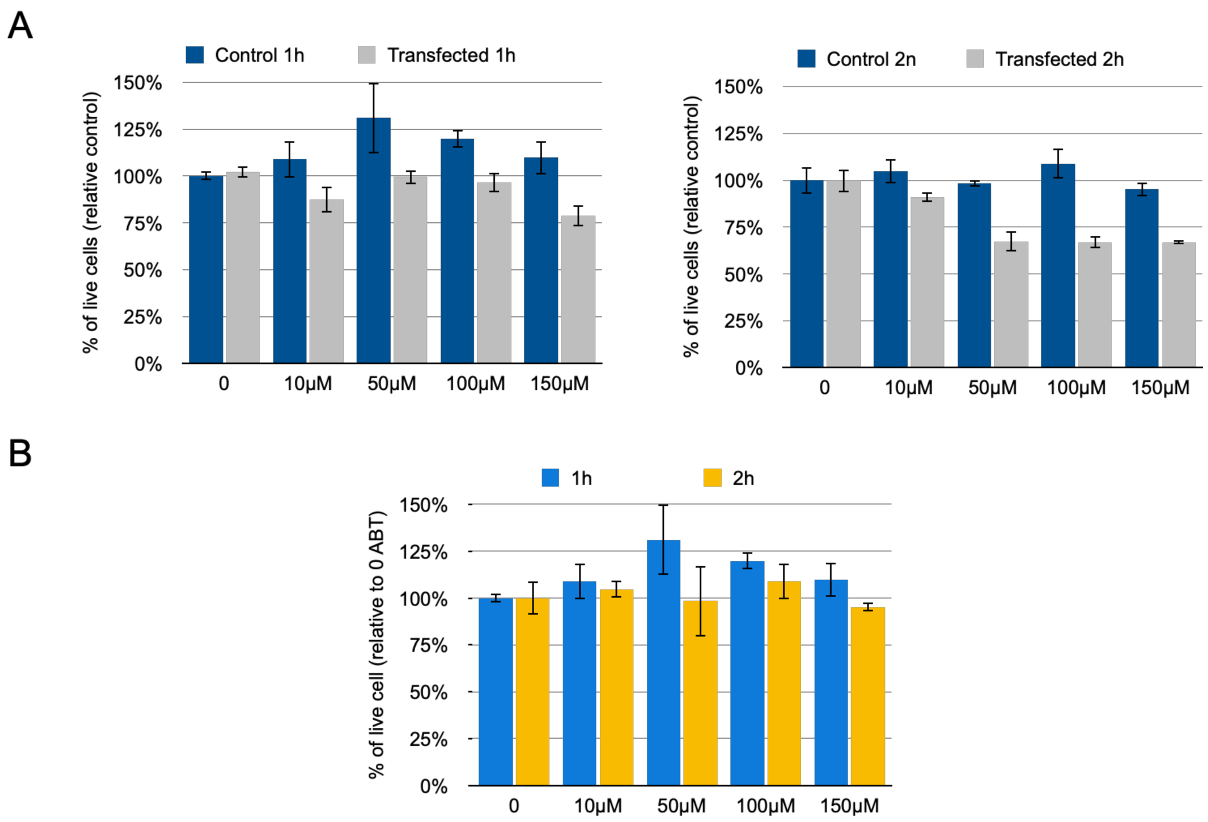


**Figure S6: HCT-116 cell viability in response to ABT-199. A.** Comparison of cell viability of HCT116 over-expressing caspase-3 to non-transfected (Control) cells to treatment with different concentrations of ABT-199. Results from 1h incubation are on the left and from 2h incubation are on the right. Cell viability was assessed by XTT assay relative to control cells treated with DMSO (0 ABT) **B.** HCT116 cells viability following 1 or 2 hours incubation with different concentrations of ABT-199. Cell viability was assessed by XTT assay relative to cells treated with DMSO (0 ABT).

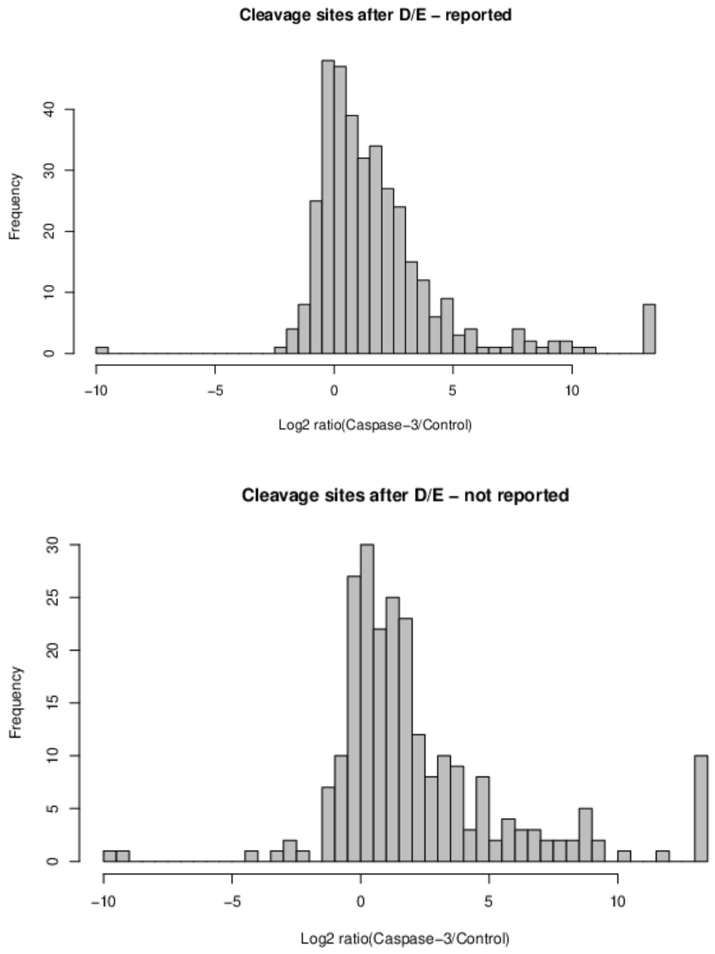


**Figure S7: Ratio distribution of caspase-3 cleavages in cell-based experiments.** Frequency distribution of caspase-3 generated peptides identified by HYTANE and LATE (combined) in HCT116 cells overexpressing caspase-3 and their control. The ratios of peptides that matched reported caspase-3 cleavage sites are on the top and those of putative unreported caspase-3 cleavage sites are on the bottom.

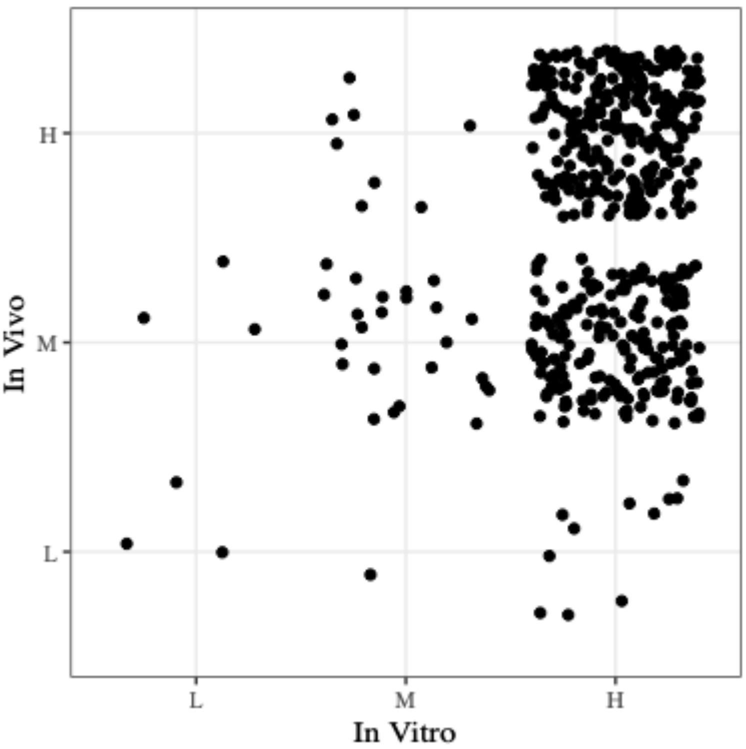


**Figure S8: Putative caspase-3 cleavages ratio comparison between *in vitro* and cell-based experiments.** The different peptides that were identified in the *in vitro* and cell-based experiments were categorised and marked into 3 groups based on the Log2 of their abundance (caspase-3/control): L= lower than -1, H= higher than +1 and M=between -1 to 1. Each peptide represented by a dot is plotted according to its ratios in both samples. Peptides that were identified in only one experiment were excluded.

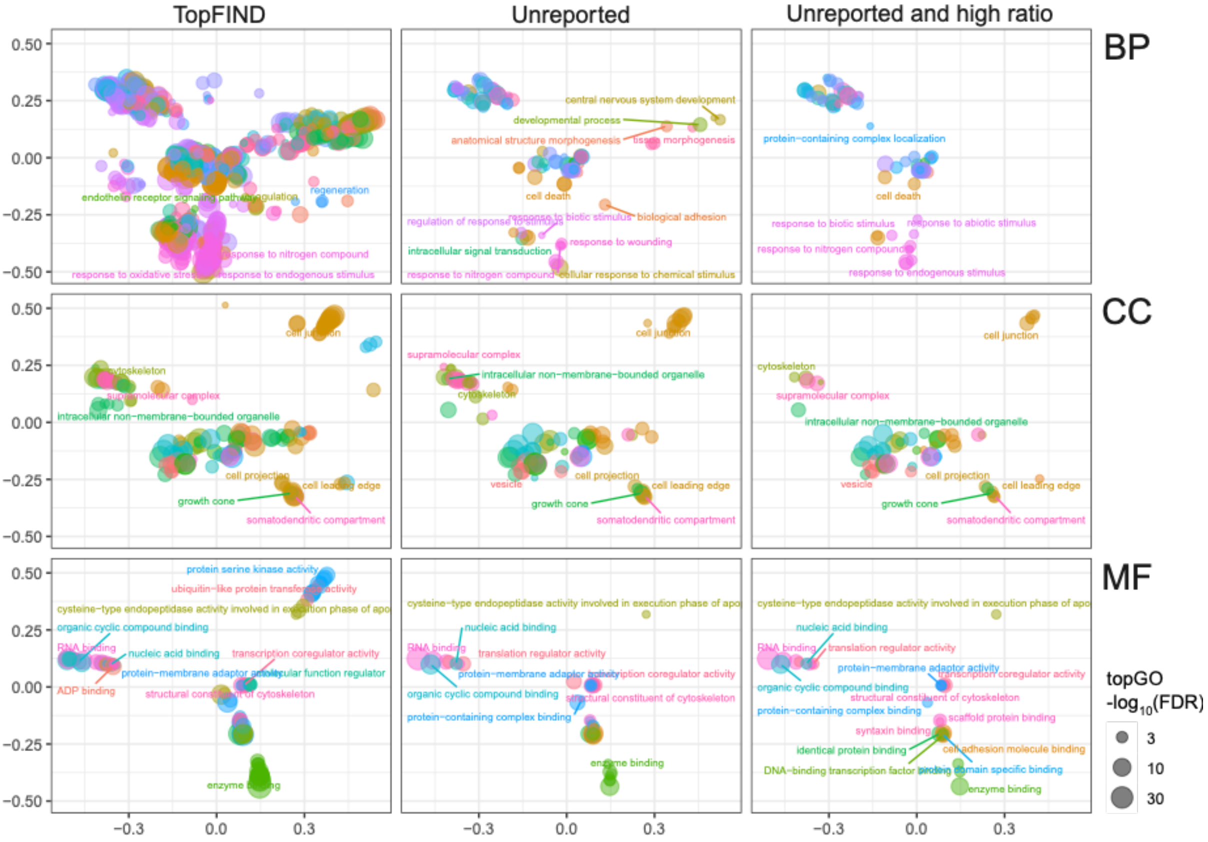


**Figure S9: Go enrichment of caspase-3 cleaved proteins.** Results of the GO enrichment analysis for proteins with caspase-3 cleavages in the in vivo experiment with the focus on the proteins not reported previously (all such proteins and those that demonstrated a high abundance ratio in the caspase-activated cells), compared to the proteins reported to be cleaved by caspase-3 in TopFIND (release 20211220). GO term enrichment analysis was performed with topGO for each category independently and those terms that demonstrated FDR q-values ≤ 0.01 were picked. Rrvgo was used to reduce redundancy and obtain semantic axes for the visualisation. See Table S12 for the numerical data used in this figure.

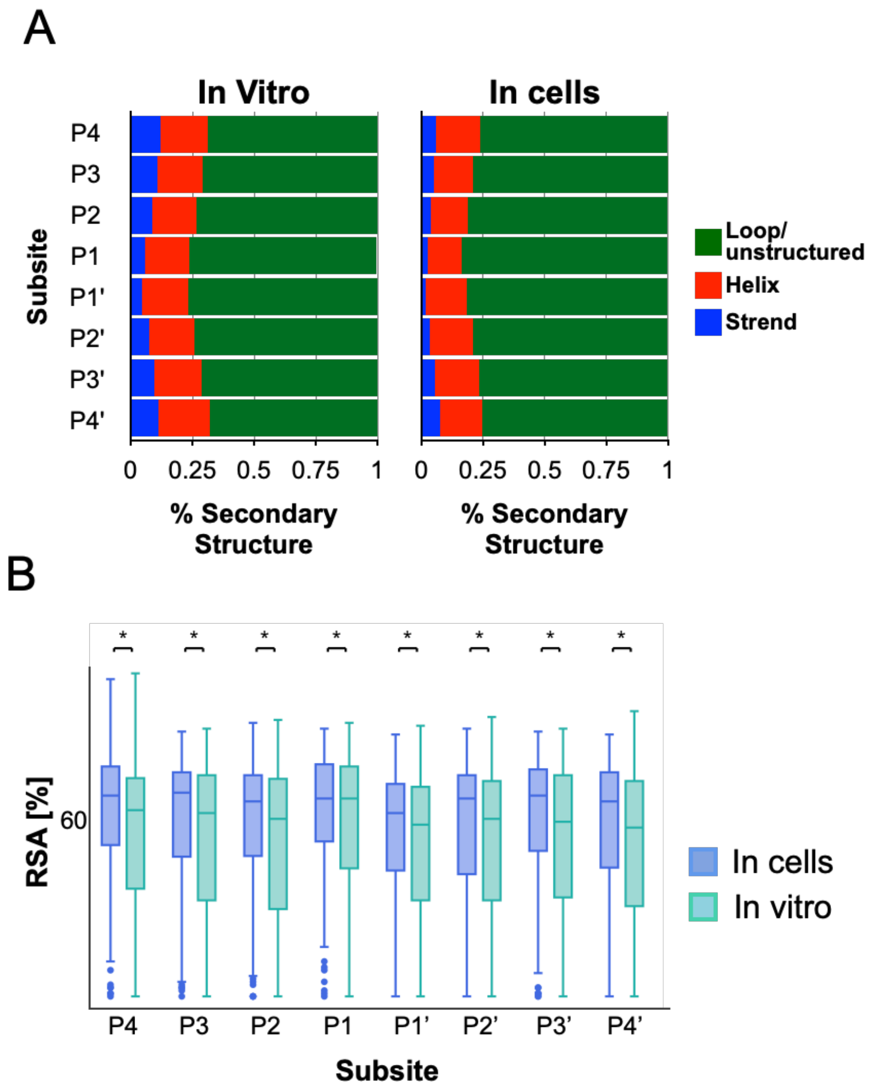


**Figure S10: Structural analysis of caspase-3 cleavage sites.** The secondary structure (A) and the relative solvent accessibility (B) of caspase-3 cleavages were investigated. The analyses include only cleavage motives (P4’ to P4) derived from cleaved peptides with Log2(caspase-3/control ratio) ≥ 1 and were identified at least twice in the cell-based experiments or the in vitro experiments. AlphaFold (Varadi et al., 2022) predictions were used to determine the secondary structure at each cleavage site and DSSP(Joosten et al., 2011; Kabsch and Sander, 1983) to determine the relative solvent accessibility (RSA). In both cases, cleavage motif secondary structure distribution is very similar when caspase-3 mostly cleaves at non-structured or loop regions, with only a small fraction of cleavages occurring at helical structures and almost no cleavages occurring at beta sheets, consistent with previous reports about this protease (Barkan et al., 2010; Timmer et al., 2009). The relative exposure of the cleavage as indicated by the RSA value changes significantly between the two experiments when the cleavages identified in cells occur in more exposed regions than those obtained in vitro. This might reflect the impact of cell lysis that is performed before the incubation with the enzyme in the in vitro experiments and can cause alteration in protein structure and cellular compartmentalization that is kept under control in the cell-based experiments.

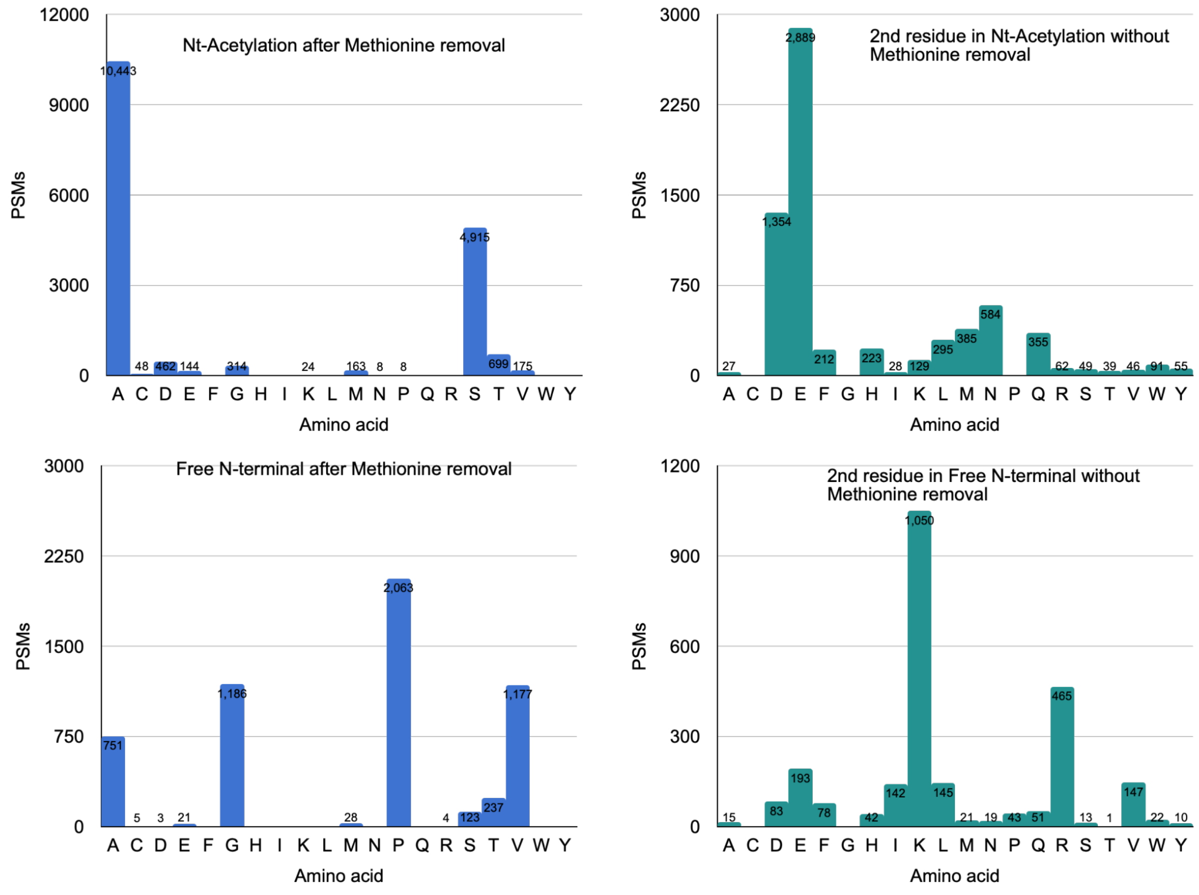


**Figure S11: Nature and frequency of amino acids at position 2 in ORF N-terminal peptides.** The frequency distribution of amino acids at the 2nd position of ORF N-terminal peptides was determined for all possible scenarios following initiator methionine removal and acetylation. The frequency distributions of peptides after initiator methionine removal are in blue and the frequency distributions of peptides without initiator methionine removal are in cyan. Top plots refer to peptides with N-terminal-acetylation and bottom plots to peptides with free N-terminal. Single-letter amino acid coding is used for amino acid occurrences.

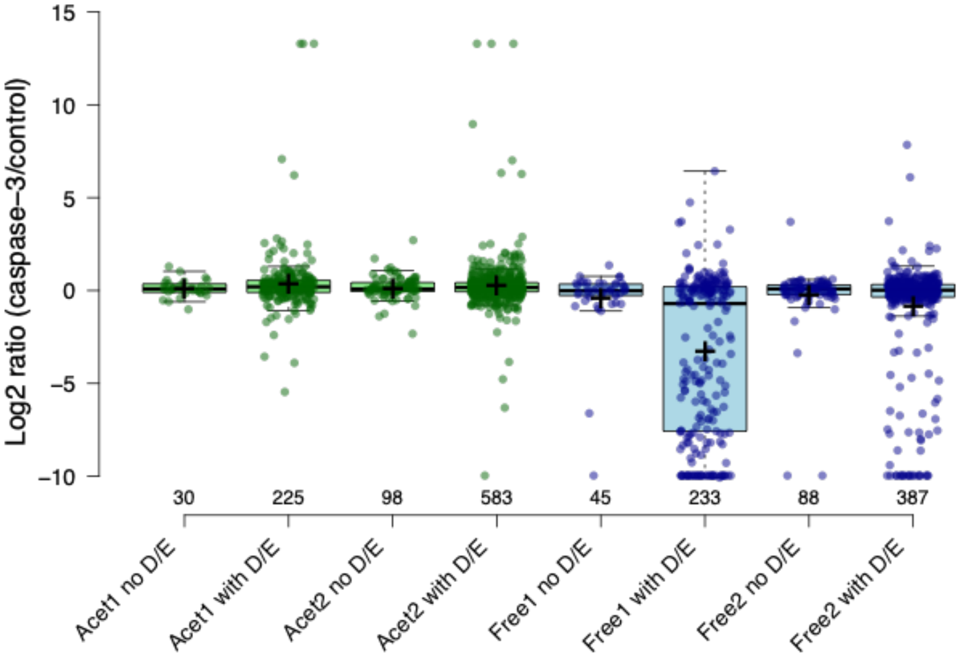


**Figure S12: The ratio distribution of ORF-N-terminal peptides as a function of initiation methionine and Nt-acetylation and the presence of Asp or Glu residues.** Identified ORF N-terminal peptides were categorized based on the presence of initiation methionine (marked by 1) or its removal (marked by 2), the presence (in green) or absence of Nt-acetylation (e.g free; in blue) and the presence of Asp or Glu residues in the peptide sequence. Center lines show the medians; box limits indicate the 25th and 75th percentiles as determined by R software; whiskers extend 1.5 times the interquartile range from the 25th and 75th percentiles, crosses represent sample means; data points are plotted as open circles. n = 30, 225, 98, 583, 45, 233, 88, 387 sample points.

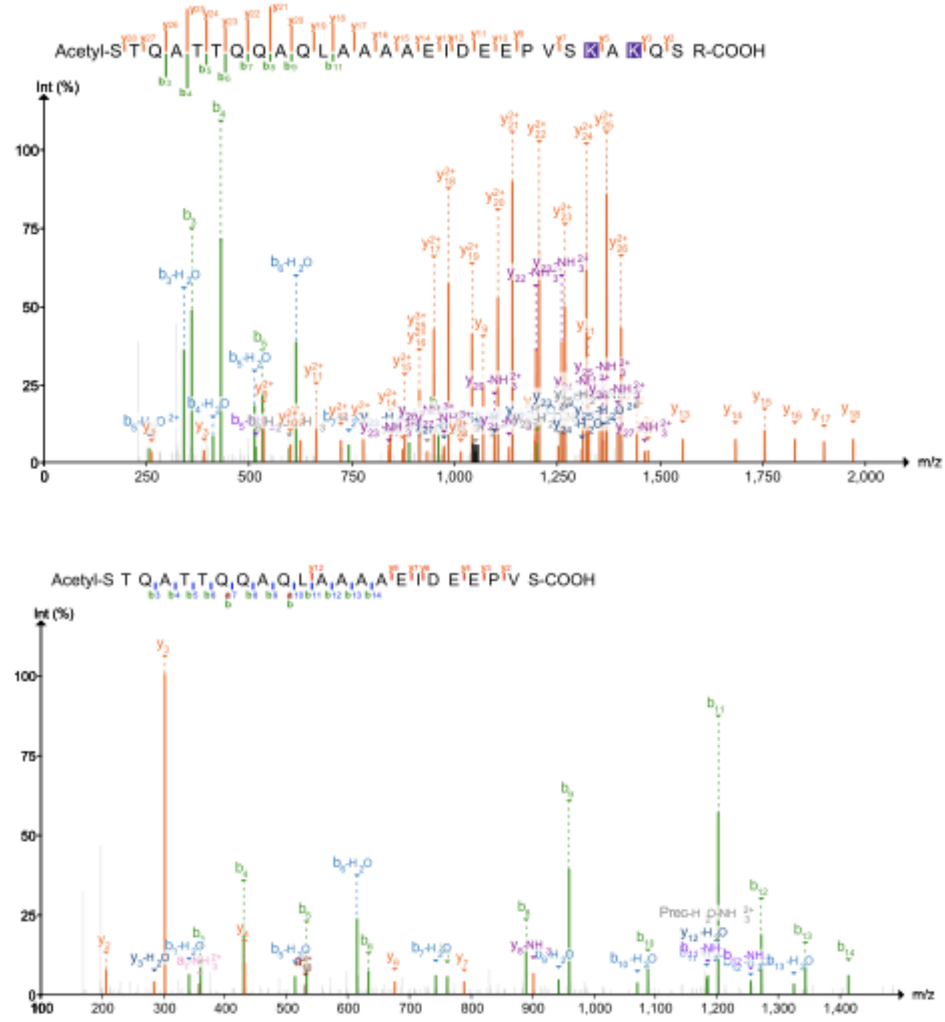


**Figure S13: MS/MS spectra of NACA Ser43 neo-Nt-Acetylated peptide**. Representative MS/MS spectra of NACA Ser43 neo Nt-acetylated identified following HYATNE (top) and LATE (Bottom). B-ions are marked in green and y-ions in orange.

| Search type | Enzyme number | Diesgtion Type | Missed cleavage | Fixed modification | Varibale modification |
|---|---|---|---|---|---|
| Hytane Nt acetylation | 5 (ArgC) | Semi | 2 | C_cysteine = 57.021464<br>K_lysine = 28.0313 | variable_mod01 = 15.9949 M 0 3 -1 0 0<br>variable_mod02 = 42.010565 n 0 1 -1 0 1<br>variable_mod03 = 6.031817 K 0 3 -1 0 0 |
| Hytane Dimethylation | 5 (ArgC) | Semi | 2 | C_cysteine = 57.021464<br>K_lysine = 28.0313<br>Nterm_peptide = 28.0313 | variable_mod01 = 15.9949 M 0 3 -1 0 0<br>variable_mod02 = 6.031817 n 0 1 -1 0 0<br>variable_mod03 = 6.031817 K 0 3 -1 0 0 |
| Hytane MetP | 5 (ArgC) | Semi | 2 | C_cysteine = 57.021464<br>K_lysine = 28.0313<br>Nterm_peptide = 14.01565 | variable_mod01 = 15.9949 M 0 3 -1 0 0<br>variable_mod02 = 3.0159085 n 0 1 -1 0 0<br>variable_mod03 = 6.031817 K 0 3 -1 0 0 |
| Hytane labelling efficency* (heavy only) | 4 (LysN) | Semi | 2 | C_cysteine = 57.021464 | variable_mod01 = 15.9949 M 0 3 -1 0 0<br>variable_mod02 = 34.063117 K 1 3 -1 0 0<br>variable_mod04 = 34.063117 n 1 1 -1 0 0<br>variable_mod04 = 34.063117 K 2 3 -1 0 0<br>variable_mod05 = 42.010565 n 2 1 -1 0 0 |
| LATE Nt acetylation | 4 (LysN) | Semi | 0 | C_cysteine = 57.021464 | variable_mod01 = 15.9949 M 0 3 -1 0 0<br>variable_mod02 = 42.01060 n 0 1 -1 0 0 |
| LATE Dimetylation | 4 (LysN) | Semi | 0 | C_cysteine = 57.021464<br>Nterm_peptide = 28.0313 | variable_mod01 = 15.9949 M 0 3 -1 0 0<br>variable_mod02 = 6.031817 K 0 3 -1 0 0 |
| LATE MetP | 4 (LysN) | Semi | 0 | C_cysteine = 57.021464<br>Nterm_peptide = 14.01565 | variable_mod01 = 15.9949 M 0 3 -1 0 0<br>variable_mod02 = 3.0159085 n 0 1 -1 0 0 |
| LATE labelling efficency* (heavy only) | 4 (LysN) | Semi | 2 | C_cysteine = 57.021464 | variable_mod01 = 15.9949 M 0 3 -1 0 0<br>variable_mod02 = 34.063117 K 1 3 -1 0 0<br>variable_mod04 = 34.063117 n 1 1 -1 0 0<br>variable_mod04 = 34.063117 K 2 3 -1 0 0<br>variable_mod05 = 42.010565 n 2 1 -1 0 0 |

**Table S11: Comet MS/MS different search parameters.** HYTANE and LATE database searches were conducted separately for each type of modification. All searches were done while defining the following parameters: isotope error - type 3, fragment bin tolerance=0.2, fragment bin offset=0 and theoretical fragment ions=0. Variable modifications are in the format of <mass> <residues> <0=variable/else binary> <max_mods_per_peptide> <term_distance> <n/c-term> <required>.

**NACA**

Casp-3 cleavage+neo-Nt-Acet

Ribosome binding

Ribosome binding inhibition

```
MPGEATETVPATEQELPQPQAETGSGTESDSDESVPELEEQDSTQATTQQAQLAAAAEIDEEPVSKAKQSRSEKKARKAMSKLGLRQVT
GVTRVTIRKSKNILFVITKPDVYKSPASDTYIVFGEAKIEDLSQQAQLAAAEKFKVQGEAVSNIQENTQTPTVQEESEEEEVDETGVEV
KDIELVMSQANVSRAKAVRALKNNSNDIVNAIMELTM
```

**NACB/BTF3**

Isoform N-term - found in our data

Ribosome binding

```
MRRTGAPAQADSRGRGRARGGCPGGEATLSQPPPRGGTRGQEPQMKETIMNQEKLAKLQAQVRIGGKGTARRKKKVVHRTATADDKKLQ
FSLKKLGVNNISGIEEVNMFTNQGTVIHFNNPKVQASLAANTFTITGHAETKQLTEMLPSILNQLGADSLTSLRRLAEALPKQSVDGKA
PLATGEDDDDEVPDLVENFDEASKNEAN
```

Casp-3 cleavage

**Figure S14: NACA and BTF3 sequences**. The sequences of NACA (Top) and BTF3 (Bottom) with indications proteolytic processing sites, neo-Nt-acetylation and ribosome binding and binding inhibitory domain. Negatively charged amino acids in the relevant regions are in red and positively charged ones are in blue.
